# Quality-preserving low-cost probabilistic 3D denoising with applications to Computed Tomography

**DOI:** 10.1101/2021.08.10.455778

**Authors:** Illia Horenko, Lukas Pospisil, Edoardo Vecchi, Steffen Albrecht, Alexander Gerber, Beate Rehbock, Albrecht Stroh, Susanne Gerber

## Abstract

We propose a pipeline for a synthetic generation of personalized Computer Tomography (CT) images, with a radiation exposure evaluation and a lifetime attributable risk (LAR) assessment. We perform a patient-specific performance evaluation for a broad range of denoising algorithms (including the most popular Deep Learning denoising approaches, wavelets-based methods, methods based on Mumford-Shah denoising etc.), focusing both on accessing the capability to reduce the patient-specific CT-induced LAR and on computational cost scalability. We introduce a parallel probabilistic Mumford-Shah denoising model (PMS), showing that it markedly-outperforms the compared common denoising methods in denoising quality and cost scaling. In particular, we show that it allows an approximately 22-fold robust patient-specific LAR reduction for infants and a 10-fold LAR reduction for adults. Using a normal laptop the proposed algorithm for PMS allows a cheap and robust (with the Multiscale Structural Similartity index > 90%) denoising of very large 2D videos and 3D images (with over 10^7^ voxels) that are subject to ultra-strong Gaussian and various non-Gaussian noises, also for Signal-to-Noise Ratios much below 1.0. The code is provided for open access.

**One-sentence summary:** Probabilisitc formulation of Mumford-Shah principle (PMS) allows a cheap quality-preserving denoising of ultra-noisy 3D images and 2D videos.

## Introduction

Computed tomography (CT) is one of the most frequently used medical imaging techniques, with over 100 million CT scans performed yearly worldwide [1]. An additional increase in the total number of CT examinations could be observed in the recent COVID-19 epidemics [2, 3]. However, distinguishing subtle CT image features relevant for diagnostics purposes typically requires significant radiation exposure and thus increases the patient’s radiation-imposed lifetime attributable risk (LAR). This, in turn, leads to an additional chance of developing a radiation-exposure attributable cancer type [1].

The quantification of the LAR is a complex challenge and requires modeling the multifactorial interplay of DNA damage and repair mechanisms, as well as incorporating random/stochastic effects that accumulate in the low-radiation regime. *In silico* simulations and analytical estimates for net effects of such stochastic radiation-triggered reactions imply a linear model for the dependence of LAR from the accumulated radiation exposure [4, 5, 6, 7] – with linear model coefficients being dependent on the patient’s age and sex, as well as on the particular type of the CT. Despite some controversy regarding the possible existence of low-radiation thresholds in the LAR models suggested by some studies [8], the linear no-threshold models (LNT) are currently recommended for LAR assessment by the committee for Biologic Effects of Ionizing Radiation (BEIR VII) of the National Academy of Sciences of the USA [5] and by the World Health Organisation [1]. Several recent epidemiological and methodological studies support the statement that a safe radiation dose does not exist [9, 10, 11, 12] and that the LAR of CT is exceptionally high for infants and children [10, 11, 12]. The approximately 14 million pediatric CT scans of head, abdomen, pelvis, chest, or spine performed each year world-wide [10, 1] would therefore lead to approximately 12.000 fatal cases of cancer, of which 4800 are attributable only to the USA.

The prognosis that the reduction of the highest 25% of doses to the median could prevent 43% of these cancers [10] naturally suggests the increased use of low- and ultra-low radiation CT (for radiation exposures down to 0.5 mGy). However, a reduction of radiation exposure results in increased image noise and thus necessitates the application of reliable image denoising and feature extraction tools. Facilitated by the rapid development of emergent machine learning (ML) and deep learning (DL) algorithms, this research on the boundary between medical radiology and informatics is attracting an increasing amount of attention over the past years [13]. The currently available CT image denoising tools can be roughly subdivided into unsupervised and supervised methods. The unsupervised approaches search for a hidden pattern without prior learning, whereas the supervised techniques aim to identify features previously learned from the training data. Unsupervised methods do not require previous training, allow high-speed computations, and belong to the most frequently-used image denoising instruments [14, 15]. They include methods based on local averaging of the data (like Gaussian, weighted Gaussian, bilateral and mean average filtering) [16, 17, 18, 14] and spectral methods (like Fourier-, wavelet- and PCA-denoising) [19, 20, 21, 22, 15]. Recent years have also seen an active development of very successful CT denoising approaches based on semi-supervised ML ideas (for example, methods based on generative adversarial networks) [23, 24] and Deep-learning algorithms for denoising- and image segmentation [25, 26, 13, 27]. The deep learning methods have been shown to be very successful for denoising and the current convention says that DL performs much better than traditional unsupervised regularized denoising algorithms.

However, recent evidences in the literature indicate that ML and DL tools can struggle when dealing with the denoising of real images, either due to the lack of an adequate training sample or to the increasing complexity (and computational cost) of the required network [28]. This is particularly true in medical imaging, where the approaches based on ML can sometimes lack accuracy [29], while DL tools tend to rely too heavily on labeled datasets and on sufficiently large training sets [30, 31, 32]. The size of the training set plays a very central role also in the denoising of CT images, where the number of instances in the training set *T* is significantly smaller than the feature space dimension *D*, corresponding to the number of voxels. A problem characterised by *D* » *T* pertains to the so-called “small-data learning challenge” [33, 34, 35, 36, 37], and represents a scenario in which ML and DL approaches are prone to quickly overfit the small training set and to achieve an unsatisfactory performance on the validation set [38, 39, 40, 41]. To tackle this issue, several alternative approaches have been proposed [42, 43], with transfer learning representing one of the most powerful alternatives [44]. Even the latter approach presents, however, some limitations that are particularly relevant in the denoising of CT images: due to the individual variation of small-scale anatomical features and of CT operation regimes, the structural similarity assumption between the source domain and the target domain is usually not fulfilled, while remains unclear the amount and type of information that needs to be transferred if we want to avoid potential drawbacks – like, e.g., negative transfer – that could actually lead to a performance worse than the starting deep learning model [45, 46]. Thus, while a combination of transfer learning and deep learning is being widely used to attempt the solution of small data problems in the denoising of medical images [47, 48, 49], the reported results can still be dissatisfactory due to the lack of efficient strategies to systematically tackle these limitations [50].

The issues described above are not the only ones arising in the small data regime characteristic for CT: a statistically-significant systematic comparison and benchmarking of the supervised learning approaches can be strongly biased by the so-called “concept drift”, i.e., a scenario in which the non-stationarity of the learning problem leads to a mismatch between the training data and the actual application data [51, 52, 53, 54, 55]. In CT imaging, such context-dependence of supervised ML and DL tools becomes particularly problematic when there is a discrepancy between the type of patient (age, sex, body size) and noise model tackled in the training set and those tested in the validation. This context-dependence and “concept drift” can quickly lead to unfair comparisons and unsatisfying performances of supervised learning methods. Last but not least, robustness of the learning methods can be strongly confined by the existence of structural constraints inherent for the ML and DL tools in the “small data challenge” regime: for example, while spectral filtering methods tend to outperform other unsupervised denoising algorithms [14], they also have a fundamental difficulty in dealing with high noise levels in the data [19, 20]. Recently, the existence of statistically-significant overfitting boundaries has been shown empirically by employing high-performance facilities: e.g., in [56], long short-term memory (LSTM) deep neural networks [57] have been shown to systematically overfit the data and to produce results which are not statistically-significant if the condition *T* ≥ 13.6*D* + 3.8 is not satisfied (where *T* is the size of data statistics and *D* is the number of features).

While regularised time series clustering approaches were recently demonstrated to operate in these “small data, large noise” regimes, even when the noise is an order of magnitude larger than the true signal [58, 59, 60, 61], these studies were only confined to one-dimensional denoising problems. A systematic comparison with a broad range of supervised and unsupervised methods is still lacking. Due to the stochastic nature of the noise in CT, a statistically-significant evaluation and comparison of different CT image denoising methods has to rely on sufficiently large amounts of CT images taken from the same patient under the same combination of controls (e.g., with the same tube current and the same tube voltage). However, obtaining such an extensive set of reference-imaging data for a particular patient without a medical necessity would be unethical. A systematic comparison of methods would additionally require combining such data for multiple patients in a sampled range of patient-specific parameters (age, sex, body size, etc.) as well as for a large number of practically-relevant combinations of CT controls. Furthermore, the standard quality measures like the Mean-Squared Error (MSE), Peak Signal-to-Noise Ratio (PSNE), and the Multi-Scale Structural Similarity Index (MS-SSIM) also rely on the availability of the reference image without noise but generated with the same set of underlying features [62, 63, 64]. Finally, combining existing CT data from different sources in a metastudy is problematic as well, due to a very high level of the individuality of the more subtle anatomic features of the human body on a small scale [65, 66] and would thus introduce a strong bias into such a comparative study, which would also lack the reference images. Furthermore, very few datasets containing CT projection data covering the low-radiation regime are currently available in open access, mainly due to the proprietary nature of this data and the (hidden) manufacturer-specific processing of the raw data [67, 68, 69, 70]. Even, when this information is available, like in the low-dose CT image and projection dataset described in [70], a systematic statistically-significant comparison is problematic since for each of the patients only a couple of images (with and without noise) are available, from overall *T* = 299 clinically performed patient CT exams - and with the radiation exposures practically not going below 3 mGy. As we will show below, this ultra-low radiation regime with radiation exposure down to 0.5 mGy and with SNR<1.0 imposes critical challenges for the bulk of currently-available denoising methods and will receive a particular attention in the tests performed below. To address these issues, we will lay two foundations in this manuscript.

**First**, we propose a pipeline for the automated patient-specific generation of synthetic CT images, radiation exposure estimatition and LAR computation, herewith following the strengthening movement in radiological research and using the synthetic images (e.g., like in the software tool CatSim) [67]. The created images are based on a data-driven estimation of CT image noise intensities and their relationship to CT control parameters[71, 66, 72, 73]. For this purpose, we combine the LNT model for the CT-induced lifetime attributable risk [5, 9, 10, 11, 12] with the data-driven models that relate CT noise variance to the CT voltage, current and the amount of radiation exposure [71, 74]. **Second**, we introduce the Probabilistic formulation of the Mumford-Shah formalism (PMS) and propose a regularized Scalable Probabilistic Approximation algorithm (rSPA) and its parallel extension DD-rSPA as new methods for denoising of 3D images, comparing their computational cost and denoising performances with the state-of-the-art methods in this field. Particular focus thereby is to investigate the possibility of reducing personalized LAR through improving the denoising performance in the ultra-low radiation regime (down to 0.1-0.5 mGy, with Signal-to-Noise Ratios below 1.0).

## Results

### Patient-specific generation of synthetic CT images, radiation exposure estimation and LAR computations

In the first step of the proposed pipeline we provide algorithms for generating synthetic noisy CT images for every relevant combination of CT control parameters, image parameters and patient dependent variables. Regarding the CT control parameters, we focus on the two most relevant ones that can be adjusted on the computer tomograph, which are the tube voltage, **kVp** and the tube current, **mA**. The CT image parameters are the standard deviation of the CT quantum noise, ***σ***, and the CT feature contrast in Hounsfield Units (HU). The patient-dependent variables for computing the overal CT-quantum noise as well as the CT-induced additional cancer risk, **r**, are the patient’s **age**, **sex**, the subject’s size, **d**, in *cm*, as well as the absorbed radiation dose density **CTDI**_**vol**_ in milligray (mGy). The initial reference data for the automated generation of a battery of synthetic test-images can be either a set of real CT-data generated using high-dose radiation (Fig. 1A), or artificially simulated data, respectively. These reference data have to be characterized by high image quality and low quantum noise (visualized in Fig. 1B), as compared to the (ultra) low-dose CT images (Fig. 1C) that naturally contain a massive amount of noise and thus result in low CT-image quality. Figure 1D gives a graphical abstract of the workflow from image generation to the subsequent comparison of the various ML/DL-denoising methods based on the accuracy of the denoised image data. Starting with high-quality reference data, a broad range of typical CT image noises is imposed in a multitude of combinations from patient-specific and CT control variables. The obtained noisy CT images are subsequently denoised using various state-of-the-art methods. Processed and denoised images are compared to the original reference data. To model the effect of noise in CT images, we deploy and compare three different alternatives: (i) an additive Gaussian noise model that was shown to provide an adequate description of quantum noise effects in real CT images on a small scale of several centimeters [71, 75]; (ii) the non-Gaussian multiplicative noise model where the quantum noise variances change with the underlying feature color; and (iii) the empirical CT noise model sampled from the real patient data.

**Figure 1:**
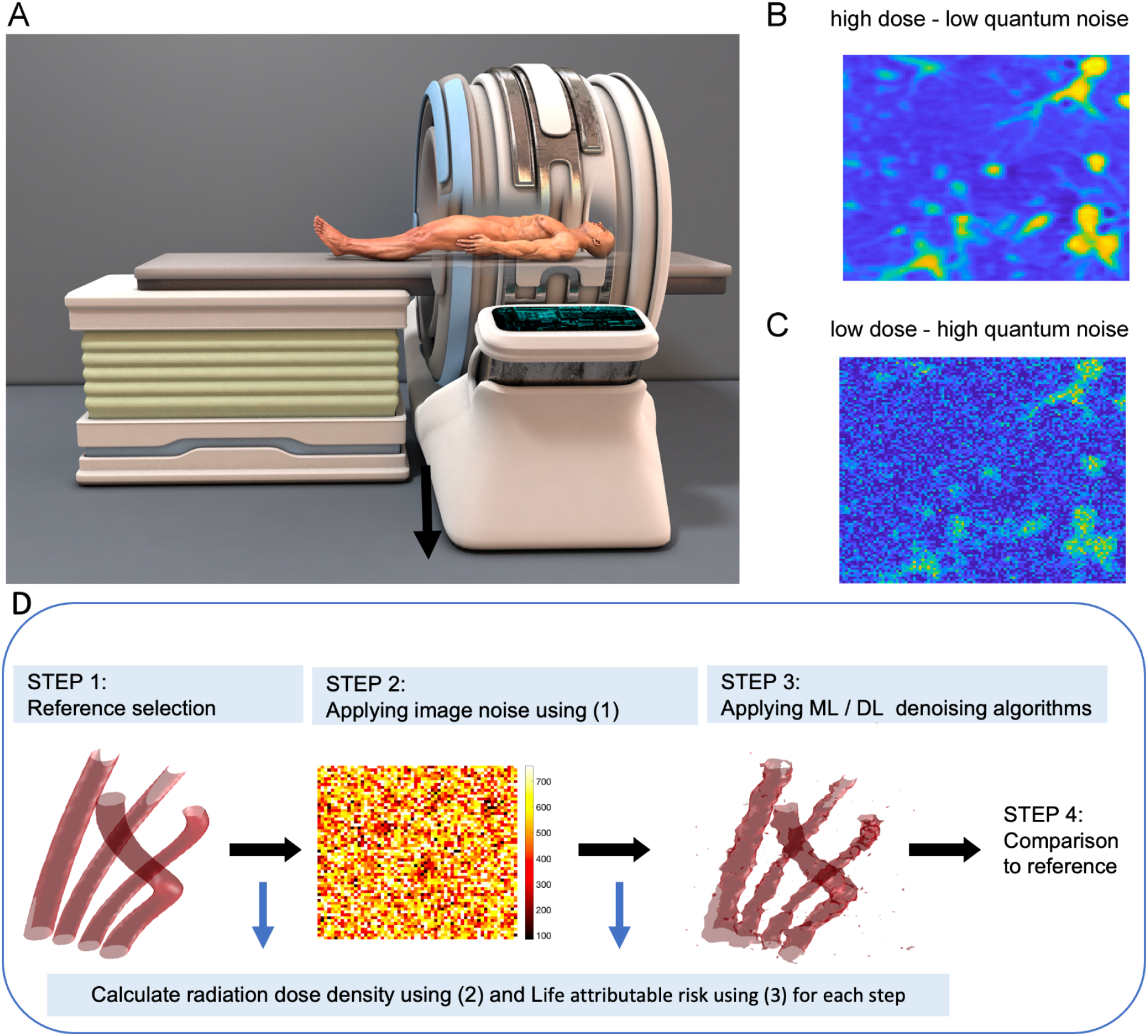
Graphical representation of our proposed pipeline’s workflow for automated generation and risk assessment of CT images. A: Initial reference data can be either a set of real CT-data generated using high-dose radiation or artificially simulated data. B: Exemplary high-quality and low-quantum noise image of lung vessels. C: exemplary low-dose CT images with high quantum noise. D: Workflow from image generation to subsequent benchmarking of ML/DL-denoising methods. Starting with high-quality data or artificially generated reference data, respectively, a spectrum of image noise ***σ*** is added for a multitude of combinations from patient-specific and CT control variables, as suggested in equation (1). The noisy images were then denoised using various state-of-the-art methods, and the processed images are compared to the original reference data.

Computation of the noise variance *σ* is performed for given CT control parameters (tube current **mA**, tube voltage **kVp**) and patient-specific parameter (water-equivalent patient diameter **d**) using the non-linear regression model introduced in [71] (see equation (1). Equation (2) of the workflow computes the effective absorbed radiation dose density **CTDI**_**vol**_ for a volume unit from the tube control parameters **mA** and **kVp** using the data-driven regression model established in [74]. Equation (3) of the image generation workflow computes the resulting lifetime attributable risk for a patient (LAR) utilizing the linear no-threshold model (LNT) proposed by the committee for Biologic Effects of Ionizing Radiation (BEIR VII) of the National Academy of Sciences of the USA [5, 76].

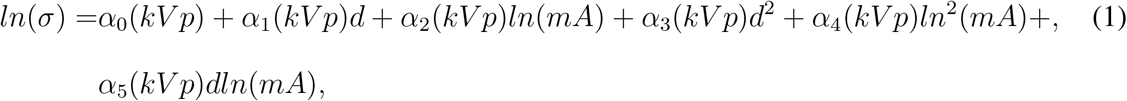

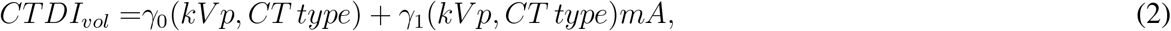

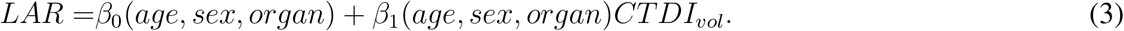

### 3D regularized Scalable Probabilistic Approximation algorithm

In the following, we introduce the 3D regularized Scalable Probabilistic Approximation algorithm (rSPA). More algorithmic details and a complete derivation with mathematical proofs can be found in the paper supplement. rSPA (see Fig. 2 for a graphical representation) seeks a simultaneous solution of image segmentation and noise elimination problems and aims to find the spatially most-persistent decomposition of the image in terms of *K* latent features. Direct application of popular segmentation and clustering methods from ML to the denoising problem results in computatinally-tractable tools with a favourable linear scaling of computational cost - but resulting in suboptimal irregular segmentations that disregard the spatial ordering of the data [77, 78, 79]. Application of regularized clustering and segmentation tools that take into account the spatial ordering and regularity of the data and features (e.g., methods based on Mumford-Shah functional optimization) have unfavourable polynomial cost scaling, limiting their application to very small images or requiring very extensive computational resources [80, 58, 59, 60, 61]. In the following, the key challenge we will address with the proposed rSPA algorithm simultaneously achieving a qualitative(in terms of low error and sufficient spatial regularity of latent features) and computationally-tractable (linearly scalable) solution of the underlying optimization problem.

**Figure 2:**
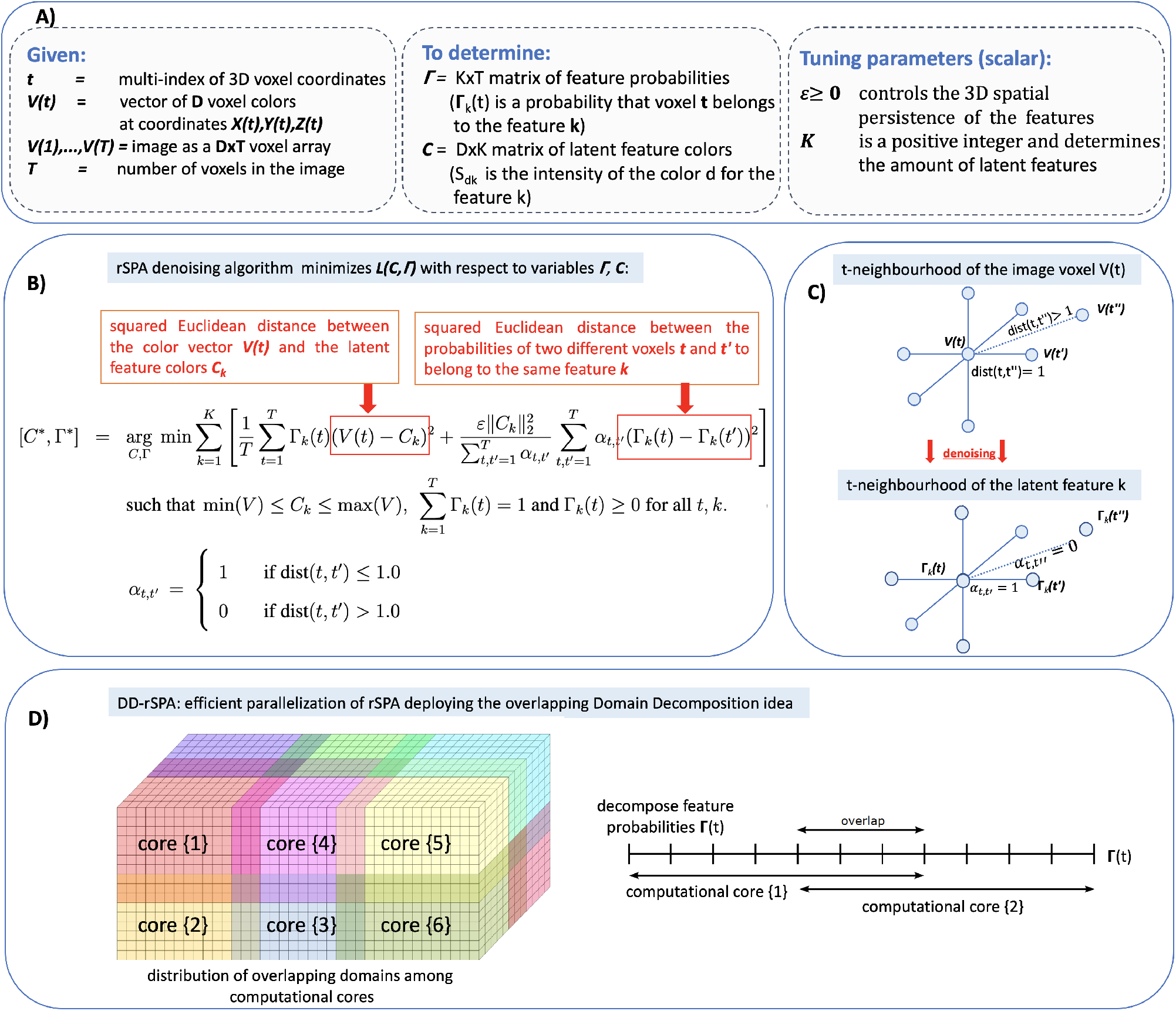
Graphical overview of the regularized Scalable Probablistic Approximation (rSPA) and its parallel extension DD-rSPA: **A)** Summary of the parameters and variables. **b)** Core rSPA algorithm idea: 3D-denoising with the regularized Scalable Probabilistic Approximation algorithm (rSPA). Given the (noisy) CT voxel data *V*, rSPA minimizes the function *L*(*C,* Γ) and seeks for the optimal segmentation of *V* in terms of the *K* spatially-persistent latent features characterized by the latent feature probabilities in *K* rows of the matrix Γ as well as by the latent colors as *K* columns of the latent color matrix *C*. Persistency of the feature segmentation is imposed by the second term of the right-hand side of the function *L*(*C,* Γ), that penalizes the differences in the feature probability values in the spatially-neighboring points. **C)**: Denoising idea: latent feature probabilities are persistent (slowly-changing) 3D functions. **D)**: graphical representation of the overlapping Domain Decomposition used in the parallel DD-rSPA algorithm.

We consider a 3D image to be provided as an array *V* = {*V* (1), *V* (2), … , *V* (*T*)} of *D*-dimensional patch value vectors for all *T* of three-dimensional CT voxels, with patch values 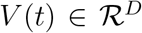 being, for example, the grey-color intensities *V*_*d*_(*t*), *d* = 1, … , *D* of the *D*-dimensional voxel patch with an index *t*. Without a loss of generality, in the following applications, we will consider the common grayscale CT images with one-voxel patches (*D* = 1) and *T* being of the order 10^5^ − 10^7^. The problem of denoising can then be considered as a numerical problem of searching for *K D*-dimensional latent features characterized by *K D*-dimensional distinct feature vectors {*C*_1,*k*_, … , *C*_*D,k*_}, with *k* taking values between 1 and *K*. Spatial characteristics of these *K* latent features we will be searching for will be provided by (a priori unknown) latent feature probabilities Γ_*k*_(*t*), representing the probabilities of an actual (noisy) voxel *V* (*t*) to belong to a particular latent (noiseless) feature with an index *k*. Such numerical procedure can be performed by a broad range of clustering and segmentation algorithms from ML (e.g., K-means, Scalable Probabilistic Approximation and others) [77, 78, 79, 81]. For example, the Scalable Probabilistic Approximation algorithm [81] would minimize the sum of the errors *L*_*t*_(*C,* Γ(*t*)) when approximating every vector *V* (*t*) with its probabilistic representation 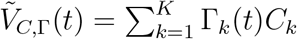:

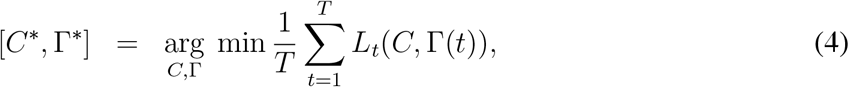

where 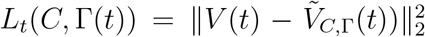. It is straightforward to see that, when *C* is fixed, the solution of the minimization problem (4) is equivalent to *T* independent minimizations of individual errors *L*_*t*_ with respect to their particular Γ(*t*) - and can be performed independently for each *t*. This allows a very efficient - independent and parallel - numerical treatment of problem (4) and results in a favourable linear scaling of the computational cost with growing size and dimension of the data [81]. The downside of this nice independent and additive structure of optimization problem (4) is that it results in solutions that are independent of any spatial permutation of the original data *V*, since the right-hand side of expression (4) is clearly invariant with respect to any arbitrary re-ordering of the summation indices *t*. This indicates that the solutions of such an optimisation problem will not change if we arbitrarily change the spatial ordering of the voxels in the original image. This invariance of the clustering outcomes with respect to the data ordering is a common characteristics of a broad class of ML methods, including, for example, Kmeans- and Fuzzy-Kmeans-clustering-methods that belong to the most popular ML algorithms, with over 3 Mio. citations according to Google-Scholar [81]. While analysing spatially-ordered data, in addition to a simple segmentation (4) of the image into *K* latent probabilistic features, we would like to enforce a spatial persistence of underlying features. To achieve this, we can enforce any two voxel points *V* (*t*) and *V* (*t*′) to have similar latent probabilities of belonging to the same features if their positions are close enough to each other. In order to deal with the relative position of the voxels, we can use the kernel function, a very popular concept in ML. The simplest alternative to measure the “closeness” of two different voxels would be provided by the Euclidean kernel, defined as a distance function *α*_*t,′*_ between two distinct points with indices *t* and *t′*:

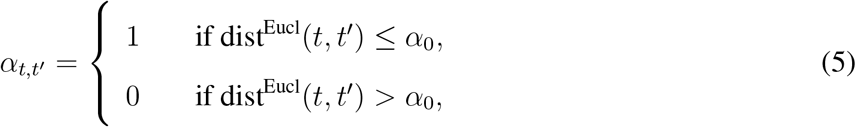

where *α*_0_ is some user-defined threshold (e.g., *α*_0_ = 1 in this paper’s applications).

Then, following the idea behind the Mumford-Shah functional formulation [80], spatially-persistent optimal probabilistic approximation 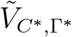 of the original image data *V* can be computed via the numerical minimization of the regularized form of the original clustering problem (4):

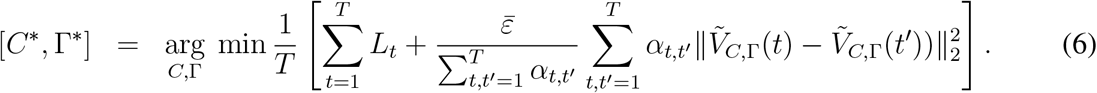

The second term in the right-hand side of this functional controls the spatial regularity and smoothness of the obtained solutions. Please note, that, in contrast to the original clustering problem (4), problem (6) is not invariant with respect to permutations of *V*, and allows to optain spatially-regular solutions [*C*\*, Γ*], with the persistence that grows when increasing the scalar control parameter 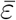. However, these nice features of the regularized problem come at a price of losing the very-favourable linear scalability of the computational cost of problem (4): optimization with respect to different Γ(*t*) can not be performed independently when *C* is fixed - as it is the case for the clustering problem like SPA (4), where one solves T independent *K*-dimensional optimization problems for Γ(*t*) with fixed *C*. The second term in (6) - that aimed at enforcing spatial regularity and persistence - at the same time introduces the global coupling between different Γ(*t*) and requires the solution of very large coupled *KT*-dimensional nonlinear optimization problems [80, 58, 59]. This confines the applicability of the image analysis methods based on (6) when working on common hardware (e.g., workstations) to relatively-small images, with *KT* not larger then 50’000-100’000 [58, 59]. Direct solution of (6) - as well as indirect Bayesian solutions of (6) based on Markov Chain Monte Carlo sampling (MCMC) - are costly beyond 1D and would require extensive use of High-Performance Computing facilities (HPC) for large realistic 3D images with *KT* ≈ 10^5^ − 10^7^ [82, 61].

One of the key methodological insights of this work is that one can systematically derive an exact upper bound approximation of the regularized problem (6) that can be solved with a linearly scalable and parallelizable numerical algorithm for realistic 3D images (with 10^6^−10^7^ voxels), while requiring few minutes on a common laptop:

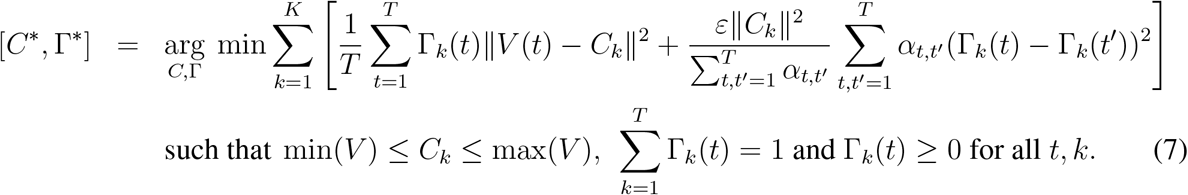

As proven in Lemma 1 of the paper supplement, solutions of problem (7) are also exact solutions of the original regularized problem (6) if the segmentations are discrete (i.e., if Γ_*k*_(*t*) take only discrete values 0 or 1). These solutions provide upper bound approximate minimizers of the problem (6) if Γ_*k*_(*t*) take fuzzy values between 0 and 1. In contrast to the original clustering SPA-functional (4), problem (7) has Γ(*t*) outside of the norm in the first (clustering) term - and the analytical structure of the second (regularizing) term is very different from the structure that one would obtain by directly deploying common regularization tools (like Ridge, Lasso and elastic net regularizations) to the original clustering problem (4)^1^

The numerical solution of the obtained optimization problem (7) can be computed with the monotonically-convergent rSPA algorithm: starting with some arbitrarily chosen *K* feature vectors *C*, one iterates solving the above problem for Γ (with fixed *C*) and minimizing of (7) for *C* (with fixed Γ). As proven in Lemma 2, 3 and in Theorem 1 of the paper supplement, rSPA always results in the monotonic minimization of (7), with a linear iteration cost scaling 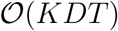. rSPA algorithm can be efficiently parallelized deploying the Domain Decomposition idea (DD) widely used in various areas. Graphical representation of the idea underlying the resulting parallel DD-rSPA algorithm is shown in the Fig. 2, a detailed description of the DD-rSPA algorithm is provided in the Section 2 of the paper supplement. Commented computer code implementing both algorithms is provided for open access at https://www.dropbox.com/sh/rw4t6ydkpi64w8y/AAA9katysG09w7ljsvUqPwwna?dl=0 and can be run on a laptop with MATLAB installation. Numerical tests on noisy images with different sizes and noise levels reveal that the overall computational cost of both the sequential rSPA and parallel DD-rSPA algorithms grows linearly withthe image size and with decreasing Signal-to-Noise ratios (corresponding to increasing noise levels), as we can see in the panel A of the Fig. 5.

### Relation of Probabilistic Mumford-Shah and rSPA algorithm to Regularized Mumford-Shah framework (MS) and Rudin-Osher-Fatemi (ROF) Total Variation model

Mumford-Shah formalism originally introduced in [80] is one of the most well-understood and elaborated theoretical and algorithmic frameworks for edge-preserving image denoising. It aims at finding the optimally-denoised image *V*^*d*^ that is simultaneously *smooth* and *close enough* to the original noisy image *V*. Then, keeping the previously introduced notation, in the most common discrete Mumford-Shah formulation such a denoised image *V*^*d*^ can be found as a solution to the following optimization problem:

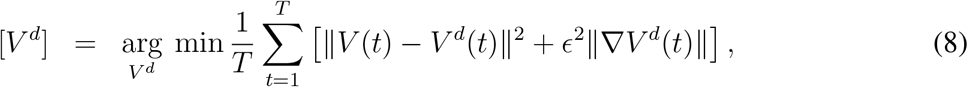

where the first term measures the “closeness” of the original and the denoised images, the second term regularizes the “smoothness” of the denoised image by penalizing the norm of its average gradient. One of the key theoretical insights to this problem (8) was provided in the work by Rudin, Osher and Fatemi [83]: deploying the Euler-Lagrange principle they shown that the solution to the minimization problem (8) is equivalent to solving a parabolic Partial Differential Equation (PDE). This opened a way of deploying the very efficient PDE solvers and the so-called *level-set methods* to the image denoising problem. The numerical solution of both the original MS-formulation (8) and of the PDE-based ROF-formalism is commonly achieved by deploying the Galerkin ansatz:

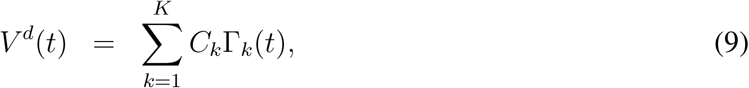

where Γ_*k*_(*t*) is a fixed set of known basis functions (e.g., mesh functions, finite element functions, wavelet basis functions, Fourier basis functions, etc.) and *C*_*k*_ are the unknown coefficients that are found numerically [83, 84, 85, 86, 87, 88].

The most important difference between the Probablistic Mumford-Shah (PMS) problem formulation (7) and the common MS- and ROF-methods is the form of the Galerkin expansion (9): (i) PMS problem (7) deploys the probabilistic expansion (9), with unknown *C* and Γ(*t*) being a priori unknown *non-parametric* probability density vectors - whereas common MS and ROF-tools dwell on a priori fixed *parametric* sets of non-probabilistic basis functions Γ. Hence, in contrast to the parametric optimization problem (8) that allows a straightforward Euler-Lagrange reformulation in form of the parabolic PDE, the introduced PMS-formulation deals with a non-parametric variational problem (7) subject to both equality and inequality constraints, that do not allow a straightforward Euler-Lagrange reformulation - and do not allow deploying the very efficient algorithms from PDE numerics for its solution. One of the central methodological developments of this manuscript was showing that despite of this presumed limitation, it is possible to efficiently solve the PMS problem numerically (7), with an iterative algorithm that has a linear scalability of computational cost. Direct numerical comparison of PMS and the common MS- and ROF-tools [83, 88] reveals very significant differences in denoising performance, cost and parallel scalability (see Fig. 5 A-C).

### Application and comparison of the rSPA method with standard methods

Next, we compare the denoising performance for a broad selection of supervised and unsupervised algorithms using the synthetic CT images generated with the above-introduced pipeline. As a noiseless CT reference we first use the patient data exemplified in Fig. 3A. It has 274’625 voxels and represents a cubic CT area of around 5*cm* × 5*cm* × 5*cm*. The data came from a high-radiation CT (180 **mA** tube current, **CTDI**_**vol**_ 15.4 mGy, section from a thorax CT of a 19 year old female patient). For each particular combination of tube-specific and patient-specific parameters, we used this reference image to create statistics of 100 different independent noisy synthetic CT images for every parameter combination.Fig. 3B shows the increase in noise when reducing radiation exposure. To illustrate the performance of DL on these data we first apply one of the most widely-used DL denoising networks: the Convolutional Neuronal Network DnCNN-3 from [25], with over 3264 citations according to Google Scholar. It was trained on a comprehensive collection of imaging datasets (including the Berkeley segmentation dataset, with over a million of image pairs for training) in a very broad range of Signal-to-Noise ratios and noise types (both Gaussian and non-Gaussian). Figures 3C and 3D show the effects of denoising by DL DnCNN-3 from [25] and rSPA, respectively in low- and ultralow-radiation CT. Fig. 3E) shows a 3-dimensional segmentation obtained from a stack of such high-radiation CT data whereas Figures 3F and 3G give the segmentation based on the images denoised using DnCNN-3 and rSPA, respectively. Figures 3E-F are all obtained from two feature isosurfaces at 625 and 200 Hounsfield Units (HU), respectively, representing the interior of blood vessels in the lung volume segment.

**Figure 3:**
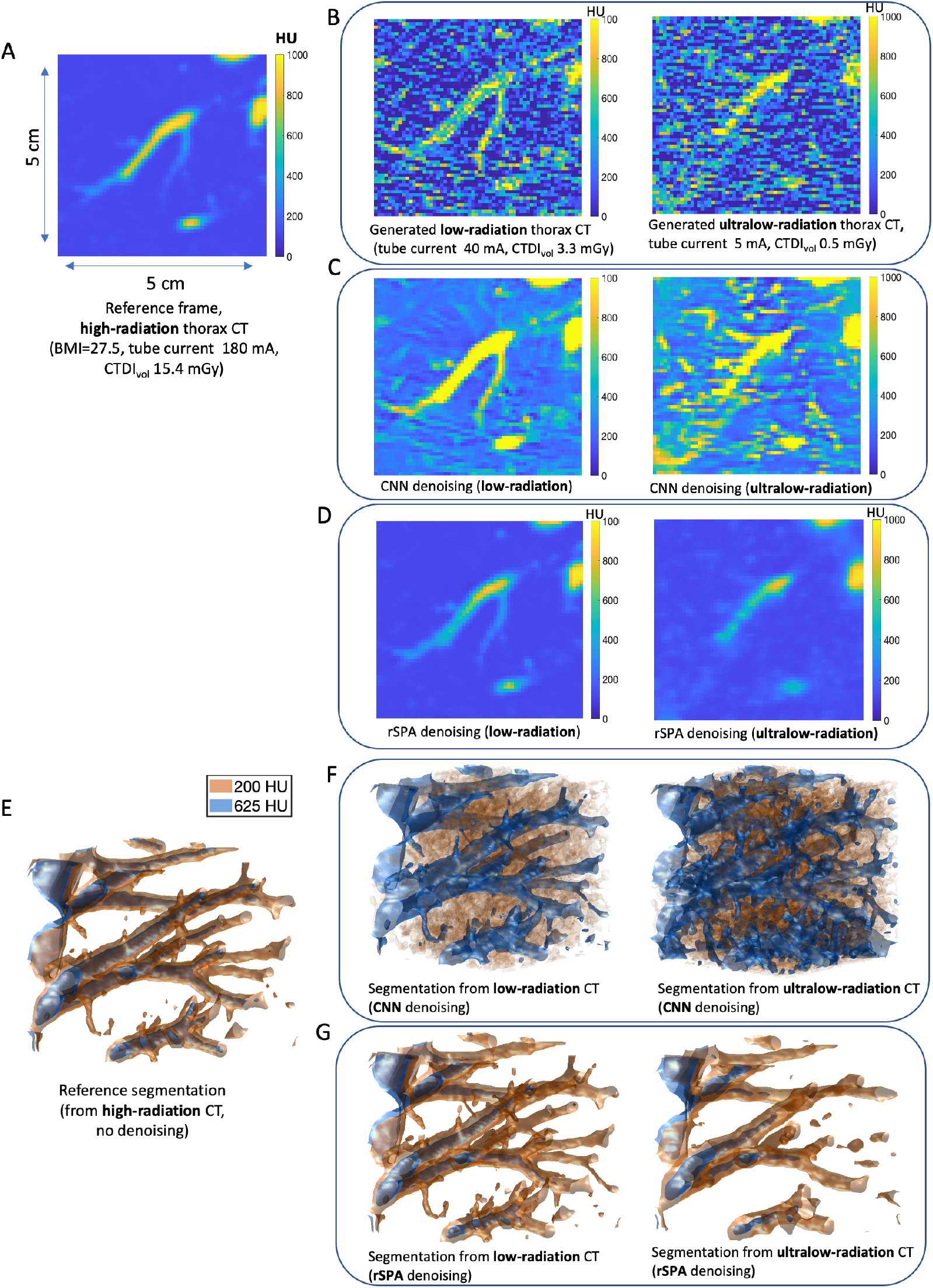
Radiation exposure, quantum noise and denoising performance of CNNs and rSPA in low-radiation and ultralow-radiation thorax CT regimes. A: Reference Data of a thorax CT voxel fragment (approx. 5*cm*^3^) of a 19 y.o. female with the BMI 27.5, acquired with the Somatum Emotion 16 2007 (Siemens) at 130 kV tube voltage. B: Simulated decreasing of the radiation exposure **CTDI**_**vol**_ from 15.6 mGy (reference frame) to 3.3mGy (for low-radiation simulations) and 0.5 mGy (ultra-low-radiation) results in a significant increase of quantum noise. C: Reconstructed images using CNNs. D: Reconstructed images using rSPA. E: 3D segmentation of the original reference frame. F: 3D segmentation based on the images denoised using CNNs. G: 3D-segmentation of the images denoised by rSPA.

Apparently, rSPA provides denoised images and segmentations that are much closer to the high-radiation reference images. Particularly, we observe that - as the noise increases - DL denoising methods start recognizing features from noise artifacts that were not part of the true reference images.

As already mentioned above, such deterioration of the performance of ML and DL methods can be attributed to various reasons, including, on one hand, the insufficient training data set and a “small data challenge” [38, 39, 40, 41] and, on the other hand, induced by the “concept drift”, stemming from the mismatch between the type of image features and the noise model used in model training and the noise model in the validation data [51, 52, 53].

To discern the potential impact of “concept drift” - and to rule-out the possibility that the “hallucinations” observed for DL CNN in Fig. 3 in the ultra-low radiation regime are induced by the insufficient training dataset - we additionally train the DnCNN-3 from [25] first with 10’000 image pairs (with and without noise) of spheres and circles of various sizes - and then with further 40’000 image pairs. We performed this two-stage training procedure to evaluate the performance improvement induced by providing more training data. The complete additional training took around 8 days on a machine with 28 CPUs (Intel Xeon Gold 6240R 2.4G, 14C/28T) and 384 GB RAM (DDR4-2933) using up to 90% of the physical cores and 120GB of memory. The resulting denoising network is provided for open access at https://www.dropbox.com/s/ia69h9fhgud2vpt/additionallytrained_DnCNN-3_network.mat?dl=0. We found that using a larger training dataset (with further 40’000 image pairs) can only bring negligible improvements, confirming the earlier finding reported in [25]. Noisy images in every pair were created using the empirically sampled non-Gaussiann CT noise at various levels, covering low and ultra-low radiation regimes (down to 0.2 mGy, corresponding to the Signal-to-Noise ratios between 5 and 0.1). In the Figure 4 we show some of the results obtained from the application of additionally trained DnCNN to the noisy images of circles and spheres that were not used in the training, deploying the same empirically-sampled non-Gaussian CT noise model as used in the training at the medium noise level (SNR=5, corresponding to the low-radiation CT) and at the high noise level (SNR=0.5, corresponding to the ultra-low radiation CT). Complete comparisons are provided as movie files and are available at https://www.dropbox.com/sh/n2dbl4h9p4o0p92/AABRkAalhXoaiKFO7ixsSzKga?dl=0. In the Fig. 4, we observe the same effect of a quick deterioration of DL denoising quality with the increasing noise as in the Fig. 3: at the medium noise level DL provides high-quality denoising, outperforming a very popular unsupervised 3D wavelets denoising tool [19, 20, 21, 22, 15]. However, at the high noise levels DL is getting outperformed by the 3D wavelet denoising. Interestingly, the best performance, in both cases, is achieved when applying the DL denoising to the data that has been previously denoised by rSPA.

**Figure 4:**
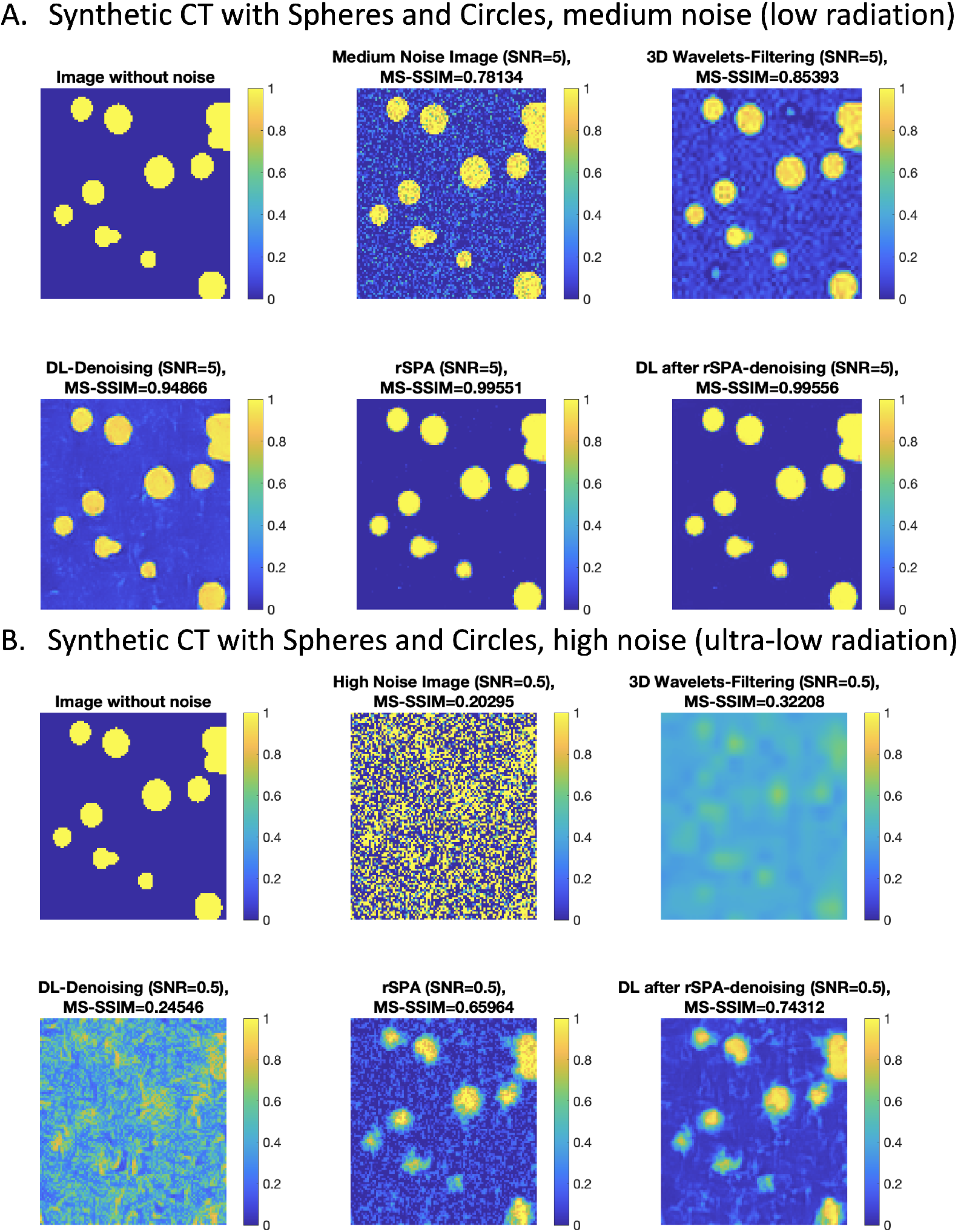
Comparing denoising performance on synthetic CT images of noisy circles, with DL from Fig. 3 additionally trained to recognize circles for non-Gaussian noise model. : A: medium noise scenario, corresponding to low-radiation regime with around 3.3 mGy; B: high noise scenario, corresponding to ultra-low radiation regime with 0.5 mGy.

**Figure 5:**
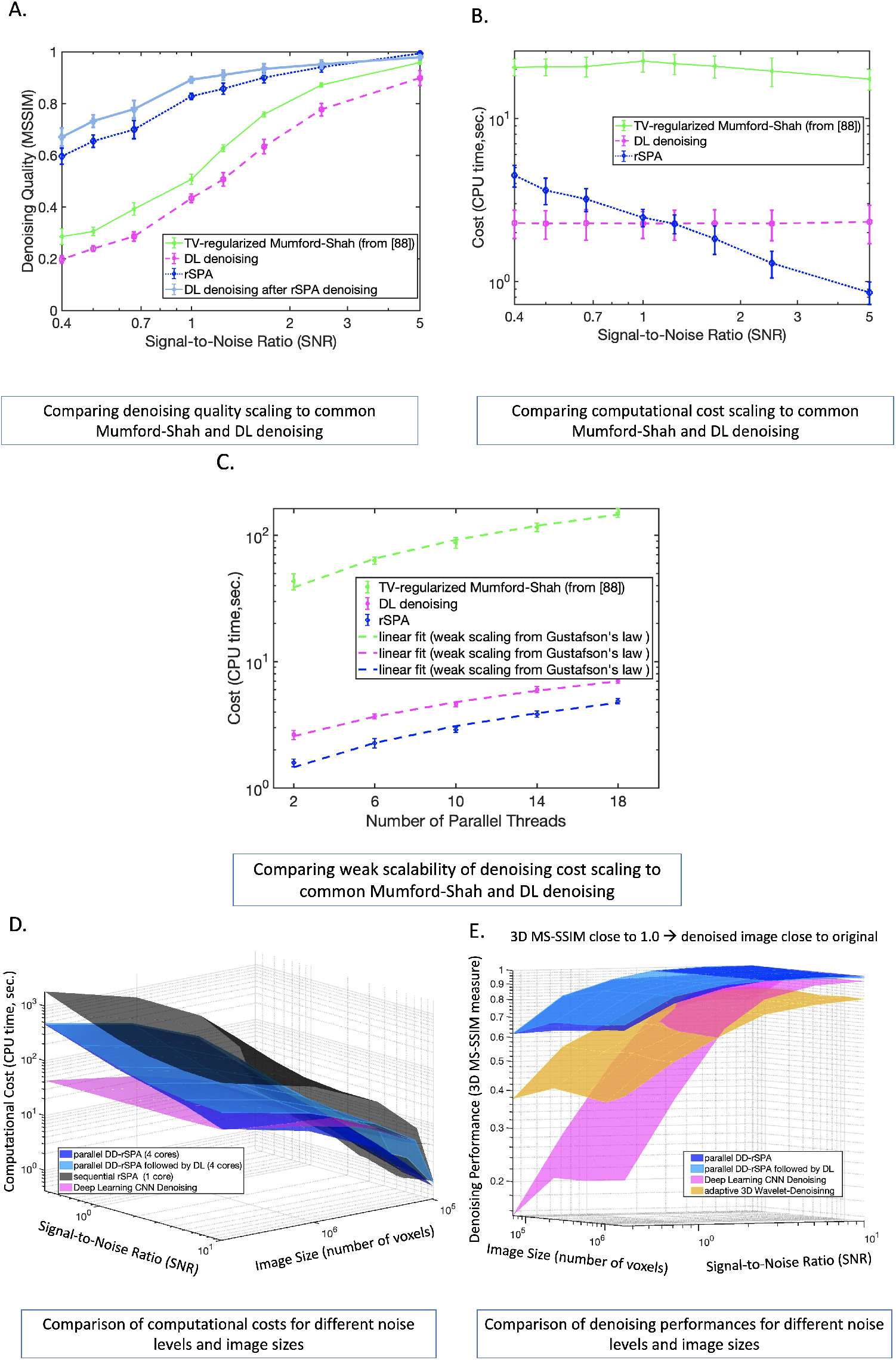
Comparing denoising quality, cost and parallelizability : A-C: comparison of PMS rSPA algorithm to the regularized Mumford-Shah denoising tool introduced in [88] and to the additionally trained DL denoising algorithm from Figs. 3 and 4; D-E: computational cost scaling and performance for DL (without taking into account time for additional training), sequential rSPA, parallel DD-rSPA and DD-rSPA followed by DL. Each point of each method’s curve and surface is obtained from statistical averaging of the respective values obtained analyzing 10 randomly-generated images with these particular combinations of image size and noise level.

Making an interim assessment of these results, we can conclude that the deteriorating performance of DL denoising is neither a result of a “concept drift” (since the type of features and the noise model deployed in the training and in validation were the same) - neither a consequence of the training data set insufficiency (since we observed only negligible performance improvements of DL when expanding the additional training data from 10K to 50K image pairs). A possible explanation can be given by the fact that here we observe a fundamental robustness boundary of DL denoising in the high noise regime, similar to the Donoho-boundary for wavelets methods [19, 20]. As we will see in the following, further numerical results provided below give additional support to this hypothesis.

In the next step, we compare the computational cost scaling, denoising performance scaling and parallelisability scalings for DL, TV-regularized Mumford-Shah denoising from [88], sequential rSPA, parallel DD-rSPA and parallel DD-rSPA followed by DL. We are particularly interested in analysing the dependence of these characterstics from the image size and noise intensity. For every combination of image size and noise level, we create 10 randomly-generated images of spheres and circles with the non-Gaussian noise - matching the characteristics of the additionally trained DnCNN-3 to avoid the bias through “concept drift”. The code reproducing these results is available at https://www.dropbox.com/sh/6p3q62zaelcyugz/AACkEjggyKcIAdgtoHGWClWPa?dl=0. The results are summarized in Fig. 5, and their computation of results required around 30 hours on a laptop with a MacBook Pro 3,1 GHz Quad-Core Intel Core i7 (4 cores) with 16 GB RAM. The measurement of the computational cost for DL considered only the pure time of applying the fully-trained DL network to a noisy image - and did not include the time needed for the additional training (that was around 8 days on the workstation as mentioned above). As it can be seen from Fig. 5, the overall costs of all considered methods scale linearly with the image size - and parallel DD-rSPA demonstrating the weak scaling of parallel computation cost (see Fig. 5C). DD-rSPA allows the denoising of a 3D image with 10^7^ − 10^8^ voxels in the ultra-low radiation regime (SNR = 0.5) at around 3-10 minutes on a MacBook Pro laptop with 4 cores. Interestingly, the costs of DL and common MS denoisings practically do not depend on the noise level, whereas the cost of rSPA and DD-rSPA grows linearly with the decreasing SNR. According to the Theorem 1 of the paper supplement, the iteration cost of rSPA and DD-rSPA does not depend on the noise intensity - and this linear dependence of the overall cost on noise is solely explained by the linear increase in the number of rSPA and DD-rSPA iterations required to achieve the solution of the minimization problem (7) with the linearly reducing SNR. In another words, these results show that DL and common MS-denoising invest the same amount of work at different noise levels, whereas rSPA and DD-rSPA invest work linearly-proportional to the SNR - and increasing with the relative increase of the noise. A comparison of the denoising quality scalings in Fig. 5 provides additional evidence towards the hypothesis formulated above: deterioration of the denoising performance of DL in the area of large noise (small SNR) and smaller image sizes - where DL is getting outperformed by the 3D Wavelet-Denoising - is not the result of an insufficient training dataset or the “concept drift”. It can be explained with the existence of a fundamental robustness boundary of DL denoising in the high noise regimes, with SNR<1.0. This finding is also confirmed by inspecting the performance of the DL when it is applied to the images that were previously denoised by DD-rSPA (light blue surface in Fig. 5B): this combination of unsupervised DD-rSPA followed by the supervised DL exhibits the best performance among all the considered methods in this high noise regime.

Next, from synthetic CT images generated from circles and spheres, we come back to the analysis of CT images generated from real anatomical features. Using the CT image generation and LAR-assessment pipeline, we compare the performance of denoising methods in a broad range of absorbed radiation dose densities. This comparison is made for two synthetic noise models (Fig. 6A and Fig. 6B, with Gaussian and non-Gaussian noise) and for the empirical nonparametric CT noise model obtained from the real patient data (Fig. 6C). The results of this comparison are shown in terms of three major image quality measures. As expected, the Gaussian 3D filtering exhibits the best performance among the common tools for all three additive Gaussian noise scenarios from Fig. 6A. On the other hand, the non-Gaussian deep learning DnCNN denoising outperforms the other tools (except rSPA) in the non-Gaussian and empirical noise situations, as can be seen in Figs. 6B and 6C. However, in the overall comparison, the rSPA method markedly outperforms all considered denoising tools in all image quality measures for all three noise models. As can be seen from Fig. 6, rSPA allows to achieve the same quality of the denoised image obtained with DnCNN (**3D MS** − **SSIM** around 0.9) with around 15-fold smaller absorbed radiation dose density (**CTDI**_**vol**_ = **0.95mGy** for rSPA vs. **CTDI**_**vol**_ = **15mGy** for DnCNN).

**Figure 6:**
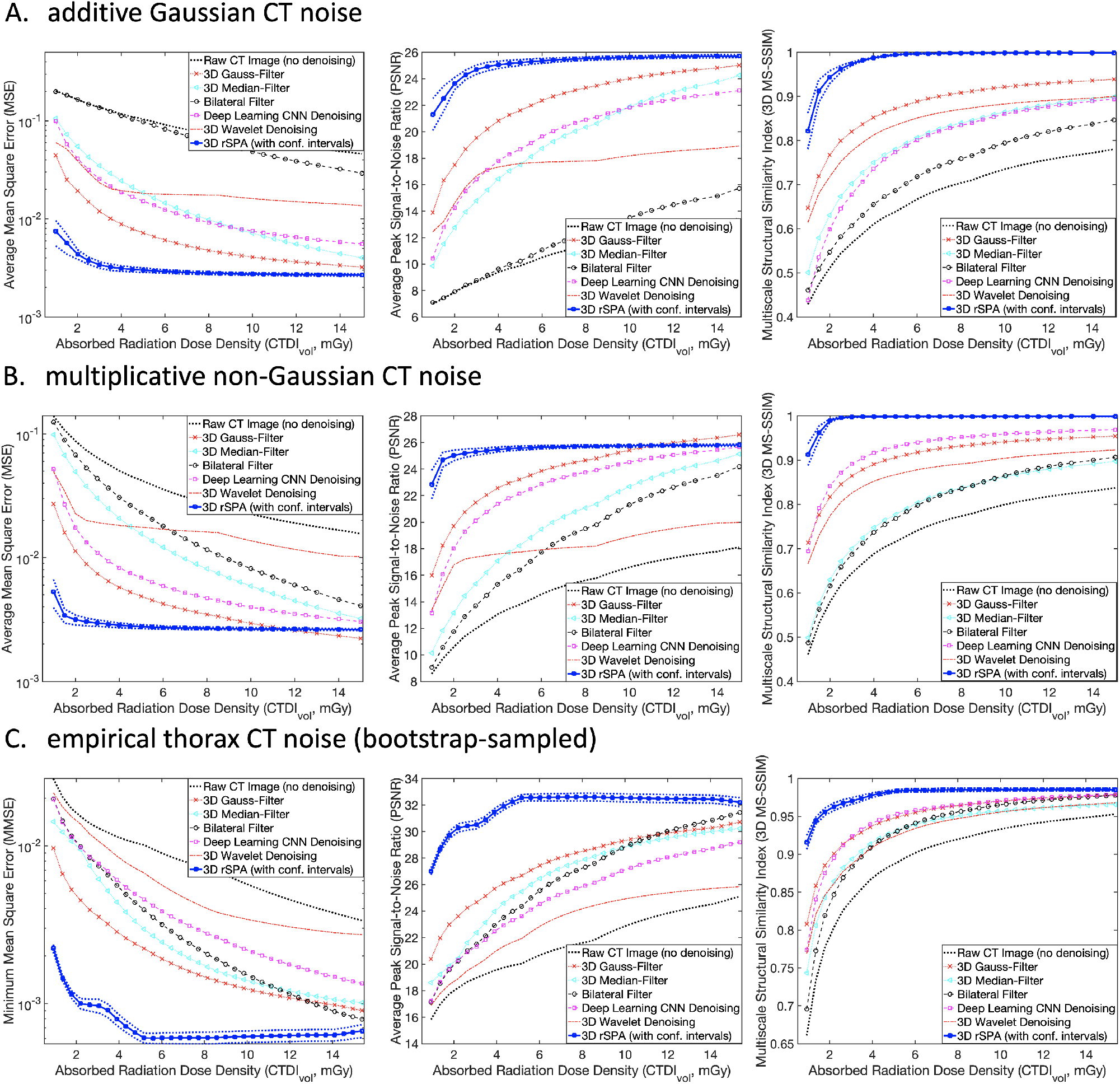
Comparing CT image denoising performances for three CT noise models. :(A) additive Gaussian noise model (CT noise variance is independent of the feature color); (B) multiplicative non-Gaussian noise model (CT noise variance changes with the amplitude of the underlying color signal); (C) empirical noise obtained from the thorax CT patient data. In (A) and (B), generation of synthetic images was performed for a patient with a water-equivalent diameter of 30 cm, which is subject to a Thorax CT with a typical tube voltage of 120 kV in the range of tube currents between 5 mA-180 mA and a set of artificial anatomic features from Fig.2A (with a feature contrast of 200 HU). In (C), real patient data were used. Comparison is performed with three primary image quality criteria: (left panels) with the mean squared error; (middle panels) with the Peak Signal-to-Noise Ratio; (right panels) with the 3D Multiscale Structural Similarity Index.

In Fig. 7, we compared the average denoising performances measured with the **3D MS** − **SSIM** image quality measure for a range of practically-relevant CT feature color intensity differences, lifetime attributable risks (LAR), and absorbed radiation dose densities. The results again demonstrate, that rSPA is superior to all other considered tools in all analyzed regimes. **3D MS** − **SSIM** of the blue surfaces corresponding to rSPA is close to 1.0 almost everywhere, indicating that the denoised images are very close to the reference CT images without noise. The powerful effect of image quality-preserving LAR reduction by denoising, especially in the female infants, is visible in Fig. Fig. 7B. Denoising with rSPA allows achieving the same imaging quality as using DnCNN (**3D MS** − **SSIM** around 0.97 for feature color differences around 50-100 HU) with a 22.6-fold smaller LAR (**LAR** = **0.015**% for rSPA vs. **LAR** = **0.34**% for DnCNN).

**Figure 7:**
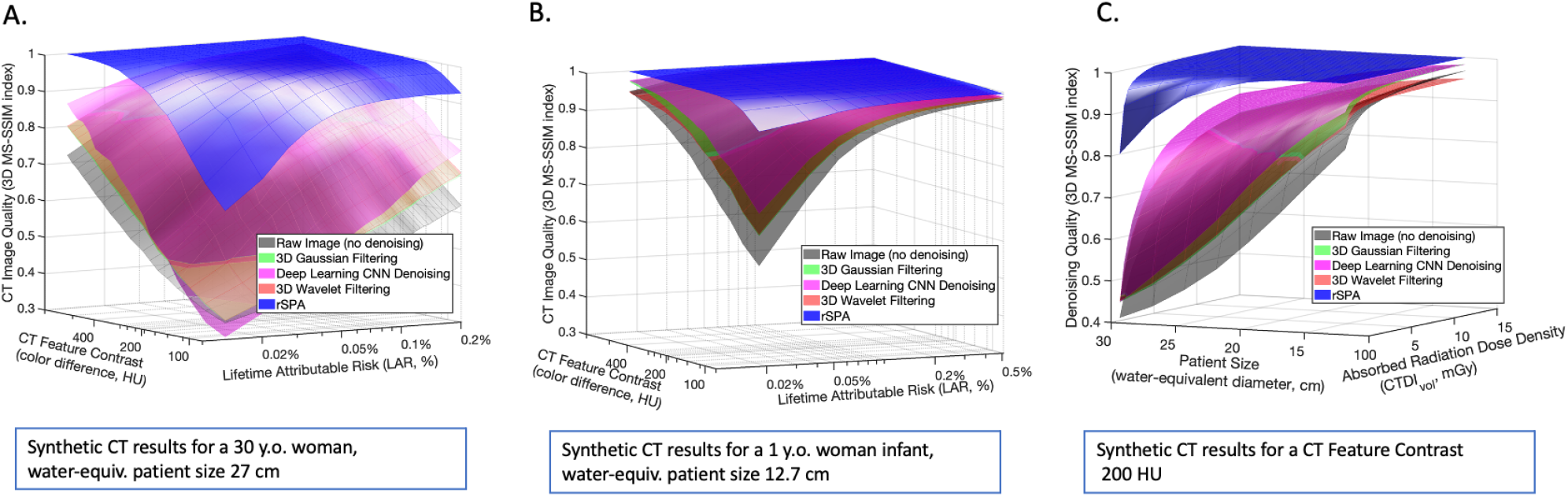
Comparing denoising methods with the average Multiscale Structural Similarity Index (3D MS-SSIM): (A) varying the true underlying feature contrast and LAR for a synthetic 30 y.o. female patient with a water-equiv. cross-section of 27cm; (B) varying the true underlying feature contrast and LAR for a synthetic 1 year old female infant patient with a water-equiv. cross-section of 12.7cm; (C) denoising performance comparison when varying the patient size and the effective absorbed radiation dose density, with the 200 Hounsfield Units (HU) feature contrast differences.

Finally, in Fig. 8 we evaluate the performance of DL with and without preliminary DD-rSPA denoising, comparing it to the denoising performance of DD-rSPA for the synthetic noisy CT images generated with real anatomic features from thorax CT. The noiseless thorax CT image used as reference in this performance comparison is available at https://www.dropbox.com/s/29x0xivg8l80q10/female_lung_thorax_CT_image_section_v2.mat?dl=0. The dotted lines show a 95% nonparametric confidence intervals (c.i.) obtained for every value of **CTDI**_**vol**_ from 100 different independently-generated noisy synthetic CT images, using the MATLAB-function *quantile()*. These results support our previous findings: applying DL to the image previously denoised with DD-rSPA provides a statistically-significant improvement of DL denoising performance.

**Figure 8:**
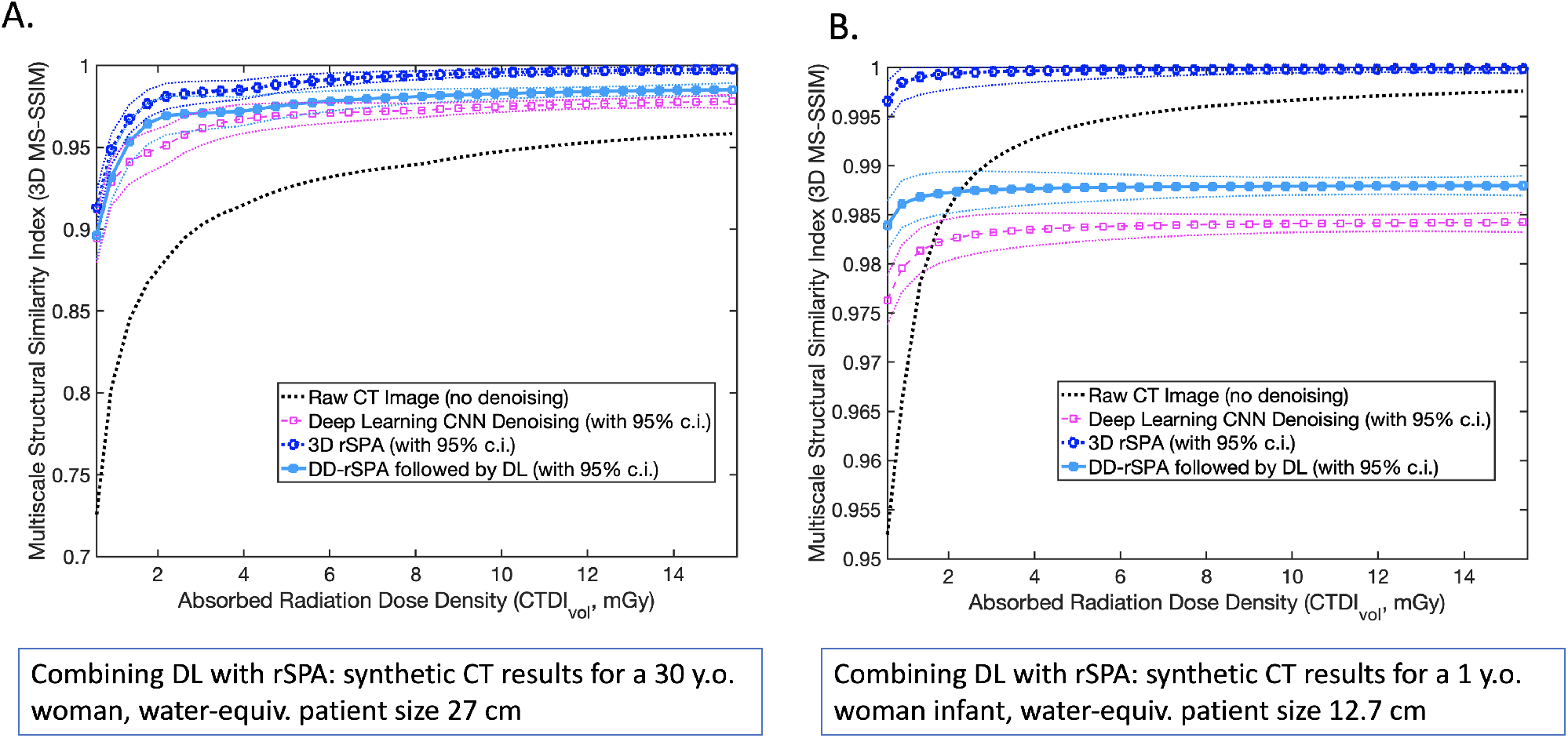
Comparing denoising methods with the average Multiscale Structural Similarity Index (3D MS-SSIM) for simulated thorax CT: (A) varying the absorbed radiation dose for a synthetic 30 y.o. female patient with a water-equiv. cross-section of 27cm; (B) varying the absorbed radiation dose for a synthetic 1 year old female infant patient with a water-equiv. cross-section of 12.7cm. Noiseless thorax CT image used as reference in this performance comparison is available at https://www.dropbox.com/s/29x0xivg8l80q10/female_lung_thorax_CT_image_section_v2.mat?dl=0. Dotted lines show 95% nonparametric confidence intervals (c.i.) obtained for every value of **CTDI**_**CTDI**_ from 100 different independently-generated noisy synthetic CT images, using the MATLAB-function *quantile()*.

## Discussion

We introduced an algorithmic pipeline for the generation of synthetic patient-specific CT images and radiation-induced risk assessment. We used it to compare various CT image denoising approaches in a range of practically-relevant CT regimes. The ultra-low radiation CT regime represents a three-fold challenge for all of the standard denoising methods: (i) reduction of the radiation exposure leads to a substantial increase of the noise, eventually making it impossible for standard unsupervised and spectral denoising tools (e.g., based on wavelets) to separate the noise from the underlying true image signals; (ii) heterogeneity and a high level of the individuality of anatomic features (e.g., of blood vessel networks) on a small scale - as well as the variability of patient sizes, CT conditions and a “small data challenge”-can lead to a problem of “concept drift” common for supervised methods, making the identification of some pre-trained features and patterns in the noisy CT images particularly difficult; (iii) as was shown in Fig. 3, in the ultra-low radiation regime performance of one of the most popular supervised denoising CNNs trained in a wide range of noise regimes [25] quickly deteriorates. To discern the potential impact of “concept drift” - and to rule-out the possible insufficiency of the training dataset - we additionally trained the DnCNN-3 from [25] first with 10’000 image pairs (with and without noise) - and then with 40’000 image pairs. We found that using a larger training dataset (with further 40’000 additional image pairs) only brought negligible improvements, thus confirming the earlier findings reported in [25].

To tackle those challenges, we introduced the Probabilistic Mumford-Shah formalism (PMS) (7) and shown that it can be efficiently solved numerically, by means of the an unsupervised regularized Scalable Probabilistic Approximation method (rSPA) that seeks a simultaneous solution of image segmentation and noise elimination problems. We could prove that it provides a computationally-cheap (with a linear cost scaling, see Fig. 5, Lemma 1–3 and Theorem 1 of the paper supplement) exact upper bound approximation of the numerically much more expensive regularized probabilistic segmentation problem (6). We also introduced DD-rSPA, a parallel extension of the rSPA algorithm based on the decomposition of the 3D-domain in overlapping subdomains (see Fig. 2 for a graphical overview, while a detailed description of the DD-rSPA algorithm is given in the Section 2 of the paper supplement). Commented code for both algorithms was provided for open access. Numerical tests with noisy images of different sizes and noise levels were summarized in the Fig. 5, revealing that : (i) the overall computational cost of both the sequential rSPA and the parallel DD-rSPA algorithms grows linearly with the image size and with the decreasing Signal-to-Noise ratios (corresponding to increasing noise levels), whereas the common Mumford-Shah and DL-denoising tools (as well as the methods like 3D wavelets denoising) “invest” the same amount of computational work independently of image SNR; (ii) the deteriorating performance of the DL denoising observed in Figs. 3,4 and 5 is neither a result of a “concept drift” (since the type of features and the noise model deployed in the training and in validation were the same) - nor a consequence of the training dataset insufficiency (since we observed only negligible performance improvements of DL when expanding the additional training data from 10K to 50K image pairs). The scaling of DL performance decay observed in Fig. 5 exhibits a much steeper robustness boundary than the Donoho-boundary [19, 20] of the wavelets denoising robustness (compare magenta and orange surfaces in Fig. 5E).

We deployed further tests that included artificial and real data, Gaussian and non-Gaussian, additive (Fig. 6A), multiplicative (Fig. 6B) and nonparametric empirical CT noise scenarios (Fig. 6C) as well as continuous and discontinuous feature boundaries. These results show that rSPA outperformed all of the other considered denoising methods in all evaluated performance measures.

The favourable linear scaling and parallelization opportunity provided by the DD-rSPA algorithm allow using a normal laptop for the tasks that would otherwise required extensive hardware (e.g., workstations and HPC facilities): as can be seen from the Fig. 5, DD-rSPA allows qualitative (with **3DMS** − **SSIM** around 0.9) denoising of a 3D image with 10^7^ voxels in ultra-low radiation regime (SNR=0.5) at around 3 minutes on a MacBook Pro Laptop with 4 cores. Non of the other denoising methods tested was able to come even close to this performance.

Results summarized in Figs. 6, 7 and 8 show that using rSPA and DD-rSPA opens a possibility to gain a significant patient-specific reduction of the radiation-imposed risks, allowing an around 20-fold estimated reduction of LAR for infants and an around 10-fold LAR reduction for adults. Based on the risk assessment protocol introduced in [10], the results from Fig.7 B indicate that adopting this personalized denoising methodology for ultra-low radiation CT in the pediatric praxis might be the key to prevent around 90% of the deadly cancers induced by pediatric CTs. This could be up to 11’000 cases yearly worldwide that can be potentially prevented.

As can be seen from Figs. 4, 5 and 8, applying DL to the images previously denoised with DD-rSPA provides a statistically-significant improvement of DL denoising. This opens a possibility to boost the performance of the supervised DL and ML methods recently developed in CT imaging. Many of the existing tools were trained in the regimes with moderate and low noise levels - and preliminary unsupervised denoising with DD-rSPA can extend their applicability to the ultra-low radiation regimes with very high noise levels.

The sequential rSPA and the parallel DD-rSPA algorithms can also be directly applied to the denoising and segmentation of ultra-noisy 2D and 3D movie data from different areas. In a case of 2D movies, time axis of a movie can be considered as the a third image dimension in rSPA. Another possible application area - the 3D movies - emerge for example in fMRI applications in various biomedical areas (e.g., in cardiology), where the main challenge is detecting the moving boundary of the inner organ and distinguishing it from other eventual shapes in a time-resolved noisy dynamics [89]. Some examples of such DD-rSPA movie denoising are available at https://www.dropbox.com/sh/n2dbl4h9p4o0p92/AABRkAalhXoaiKFO7ixsSzKga?dl=0. Finally, beyond CT data denoising and segmentation, we also see direct application possibilities for other imaging techniques, such as: fiber-optic fluorescence imaging, diffusion tensor imaging and for large-scale 3D segmentation tasks from electron microscopy images.

## Methods

### Synthetic CT image generation model

To create the additive Gaussian CT noise, we used the parameter value *’gaussian’*, non-Gaussian multiplicative noise images were created using the function *imnoise*() with the parameter value ‘speckle’ The variants parameter *σ* is in both cases selected according to the description below. MATLAB code implementing this CT image generation workflow is available at https://www.dropbox.com/sh/rr0no9vdo8osx44/AAAHQxXJnxT8P0LPs7wTRBv7a?dl=0. Generation of the nonparametric empirical CT noise was implemented in the function *create*_*CT* _*image*_*noise*() available at https://www.dropbox.com/s/xbwwrk9y2napgpy/create_CT_image_noise.m?dl=0.

### Common CT image denoising and image quality assessment methods

We used the same software platform (MATLAB) and the same hardware (Mac workstation with 28 CPU cores) for all calculations to guarantee a fair comparison of the denoising methods and to rule-out the software- and platform-induced differences that can bias this comparison. All deployed common denoising and image quality assessment tools are available in the MATLAB functions from the “Image Processing”, “Deep Learning”, “Machine Learning” and “Wavelets” toolboxes of MathWorks. We used denoising methods based on local window filtering of the data (3D Gaussian filtering with the MATLAB-function *imgaussfilt*3(), 3D local median filtering with the MATLAB-function *medf ilt*3() and bilateral filtering with the MATLAB-function *imbilatfilt*()) [16, 17, 18, 14], spectral denoising methods (the 3D wavelets denoising with the MATLAB-function *wavedec*3()) [19, 20, 21, 22, 15] and a deep learning denoising method based on pre-trained feed-forward denoising convolutional neural networks (DnCNNs, with the MATLAB-functions *denoiseImage*() and *denoisingNetwork*()) [25, 26, 13, 27]. For each of the considered images, the standard deviation of the local Gaussian smoothing kernel *σ* was changed in the range *σ* = [0.2, 0.4, 0.6, *… ,* 2]. The value leading to the least MSE deviation between the denoised and the original CT image was taken to compute the curves in Fig. 4 and Fig.5. Similarly, for the optimal 3D wavelet filtering all of the wavelet bases available in MATLAB were checked for all of the possible depths of level decompositions - and the wavelet decomposition with the minimal MSE error was taken. Pre-training of DnCNN was done with over 20 Mio images and was provided in the “Deep Learning Toolbox”. Image quality measures plotted in Fig. 4 and Fig. 5 were computed using the MATLAB-functions from the “Image Processing Toolbox”: 3D mean-squared error (MSE) [62] was computed as the average over the 2D MSE errors obtained with the MATLAB-function *immse*(); 3D Peak Signal-to-Noise Ration (PSNR) was obtained as an average over the 2D PSNR image error measures [63] implemented in the MATLAB-function *psnr*(); 3D Multi-Scale Structural Similarity Index Measure (3D MS-SSIM) [64] with the 3D image volume measure MATLAB-function *multissim*3().

### Statistical post-processing

The curves in Figures 6–8 show averages over individual denoising results obtained for 100 different independently-generated noisy synthetic CT images that were obtained for every particular combination of tube-specific and patient specific parameters. In Figure 5, the surfaces represent averages over 10 randomly-realized noisy CT images. To provide a fair comparison, same random CT image realizations were used with every denoising method. Dotted lines in Figures 6 and 8 show 95% nonparametric confidence intervals (c.i.) computed with the MATLAB-function *quantile()*.

### Data and code availability

Code is available for open access at https://www.dropbox.com/sh/rw4t6ydkpi64w8y/AAA9katysG09w7ljsvUqPwwna?dl=0 under the BSD 3-Clause License.

## Acknowledgement

The authors would like to thank Piotr Didyk (USI Lugano) for helpful discussions and comments. This work was supported by the “Emergent AI Center” of the JGU Mainz (financed by the Carl-Zeiss-Stiftung) and by the Mercator Fellowship in the DFG Collaborative Research Center 1114 “Scaling Cascades in Complex Systems”.

## Supplementary material

This document provides supplementary material for the manuscript entitled: Order of magnitude risk reduction in Computed Tomography with the unsupervised machine learning denoising. In particular, we provide the complete mathematical formulation of 3D regularized Scalable Probabilistic Approximation (rSPA) optimization problem and present:

- **Lemma 1** - derivation of the rSPA problem formulation,
- **Algorithm 1** - a subspace algorithm for solving rSPA optimization problem in pseudo-code. The algorithm consists of two consequent inner optimization problems; namely so-called *C*-problem and Γ-problem.
- **Lemma 2** - the solvability and the computational cost of solving the *C*-problem,
- **Lemma 3** - the solvability and the computational cost of solving the Γ-problem,
- **Theorem 1** - the properties and the computational cost of solving rSPA problem. The proof is based on Lemma 2 and Lemma 3.

### 1 Regularized Scalable Probabilistic Approximation Algorithm (rSPA)

**Formulation:** Let *t* ∈ ℕ^3^ be a multi-index of 3D voxel coordinates and

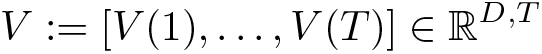

be a 3D CT image represented as a matrix of given *D*-dimensional voxel colours at 3D coordinates *X* ≔ [*X*(1), … , *X*(*T*)] ∈ ℝ^3,*T*^.

We will be searching for a probabilistic approximation 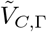 of the image in terms of *K* latent features characterized by *K* distinct color vectors {*C*_1,*k*_, … , *C*_*D,k*_, with *k* taking values between 1 and *K*. Spatial characteristics of these *K* latent features that we will be searching for will be provided by (a priori unknown) latent feature probabilities Γ_*k*_(*t*), being the probabilities of an actual (noisy) voxel *V* (*t*) to belong to a particular latent (noiseless) feature with an index *k*:

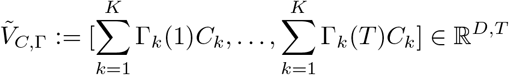

Then, following the idea behind the Mumford-Shah functional formulation [6], spatially-persistent optimal probabilistic approximation 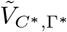 of the original image data *V* can be computed via the numerical minimization of the function:

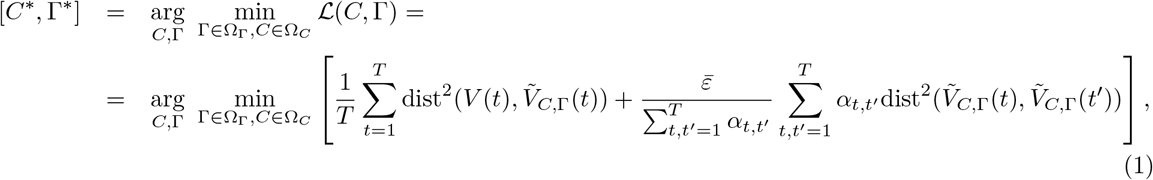

where dist^2^(·,·) is a square of some distance (e.g., Euclidean distance, *l*_1_-distance, etc.), feasible sets are given as

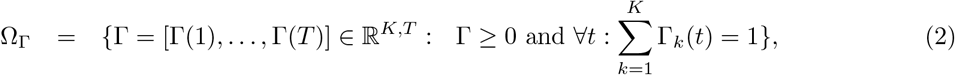

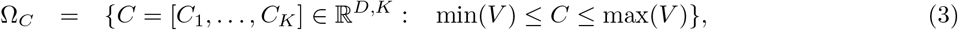

and function *α*_*t,t′*_ is the indicator function of the voxel neighborhood defined as

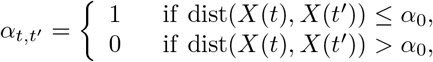

with 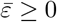 and *α*_0_ > 0 being some user-defined parameters.

#### 1. Lemma

*(rSPA as an approximate upper bound formulation for probabilistic segmentations with Euclidean distance) Approximate solutions of the problem (1,2,3) with Eulidean distance*

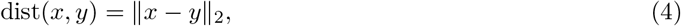

*can be found minimizing its upper bound*

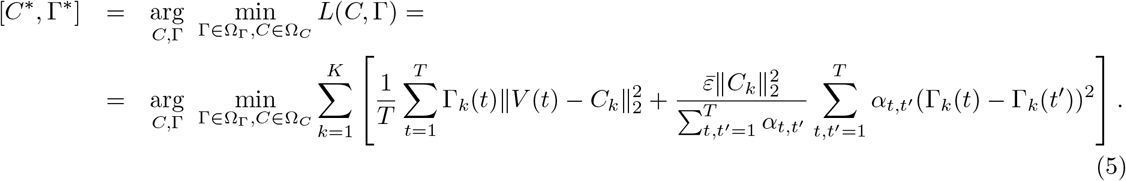

Moreover, 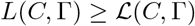 ( *for all C,* Γ ).

*Proof.* Since the square of any norm is a convex function and for any *t* = 1, … , *T* coefficients Γ_*k*_(*t*) forms the coefficients of convex combination, we can apply Jensens inequality to the first term of (1) and obtain

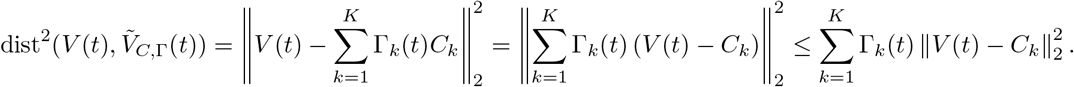

In the case of the second term, we use the properties of the norm

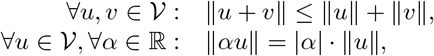

and we get

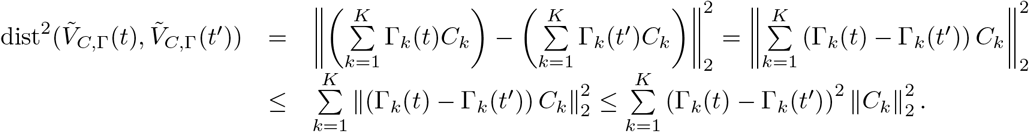

**Algorithm:** Approximate solutions of the optimization problem (5,2,3) can be found using the iterative subspace algorithm, i.e., it is solved as a sequence of split optimization problems, see Algorithm 1.

#### Algorithm 1: Regularized Scalable Probabilistic Approximation algorithm (rSPA).

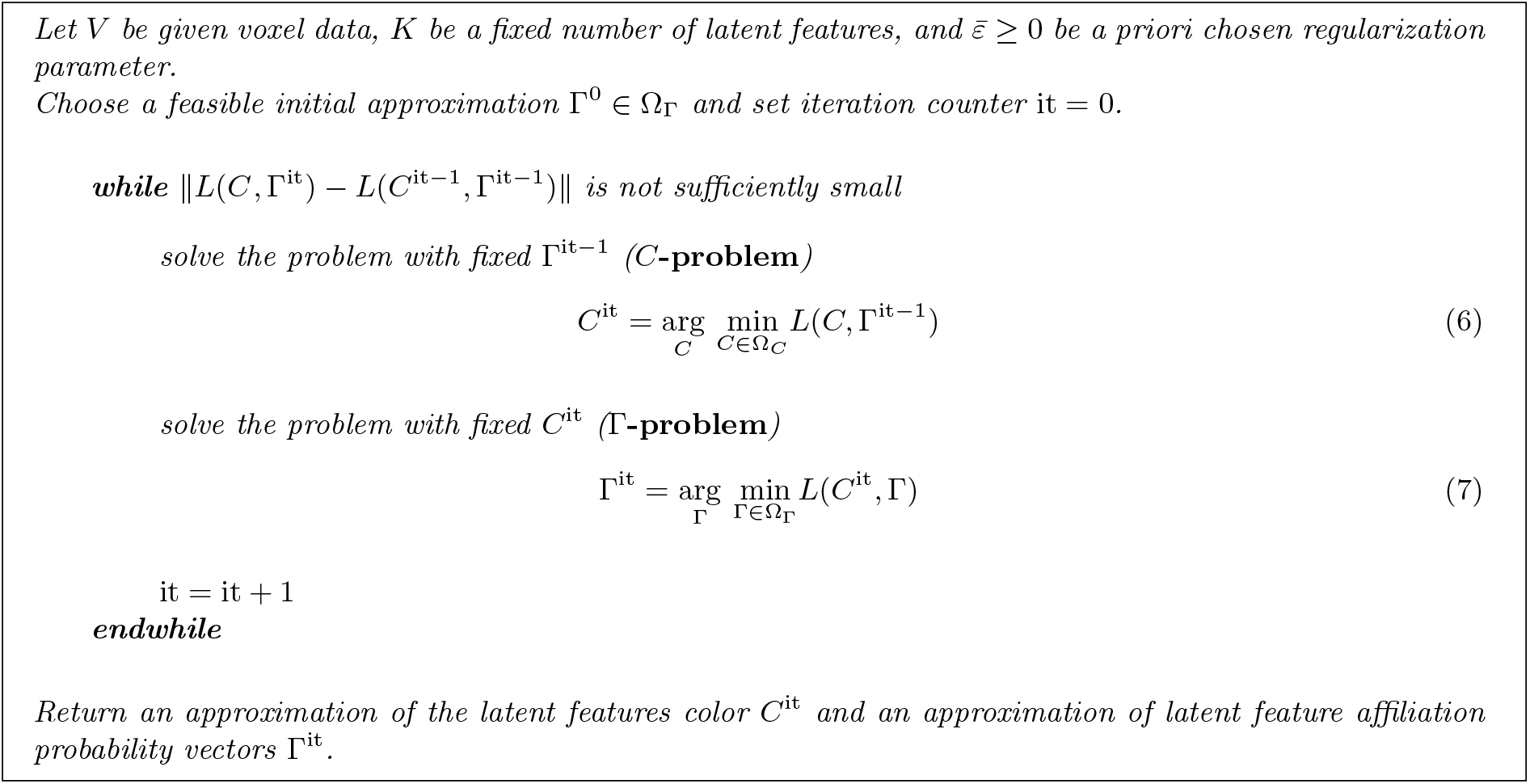

#### 2. Lemma

*(The properties of C-problem* (6))

1. *the optimization problem* (6) *has always solution,*
2. (6) *is a box-constrained convex Quadratic Programming problem (QP) with diagonal Hessian matrix and it has analytical solution,*
3. *evaluation of analytical solution for solving problem* (6) *is* 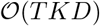.

*Proof.*

1. Let 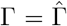 be fixed. We are dealing with minimization problem with continuous convex objective function on closed set, therefore by Weierstrass Extreme value theorem [3], the problem has always solution.
2. The objective function of problem (5) with Euclidean measure (4) is given by

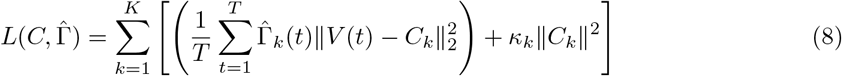

where we denoted constant

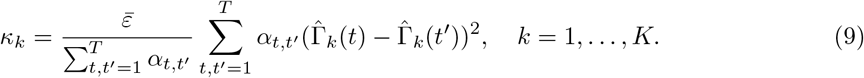 Since the minimization of (8) with respect to (3) is separable in *k* = 1, … , *K*, we have

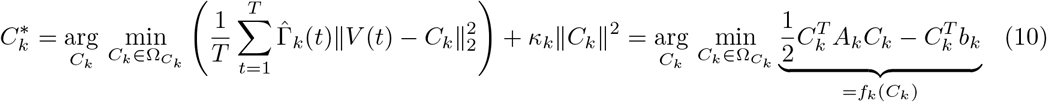

with

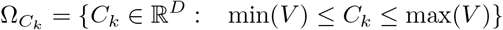

and

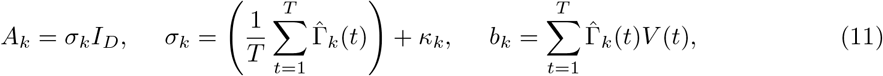

where *I*_*D*_ ∈ ℝ^*D*^ is identity matrix. Please, notice that for any non-empty cluster (i.e., 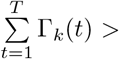 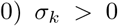 and therefore (10) is stricly convex optimization problem on closed convex set and consequently (10) has unique solution. If the cluster is empty, then (10) can be simplified to

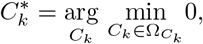

which has infinite number of solutions, i.e., any 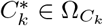 solves the problem. The problem (10) is (again) separable in *d* = 1, … , *D* and we can write

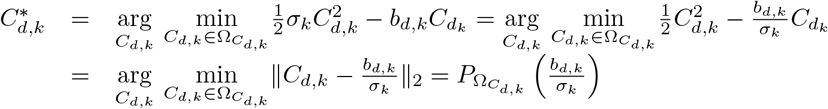

with interval

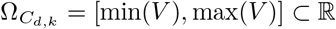

and projection onto this interval 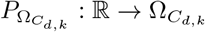 given by

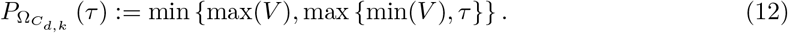
3. The computation of *K* sums of *T* vectors of dimension *D* in (11) and (9) followed by the computation of projection (12) is 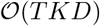.

#### 3. Lemma

*(The properties of* Γ*-problem* (7))

1. *Problem* (7) *is a convex QP on separable simplexes (i.e., with bound inequality and linear equality constraints).*
2. *Assembling the objects of problem* (7) *is* 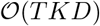.
3. *One iteration of Spectral Projected Gradient method for QP (SPG-QP, [7]) for solving problem* (7) *is* 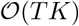.

*Proof.*

1. Let 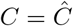 be fixed. The objective function of problem (5) with Euclidean measure (4) is given by

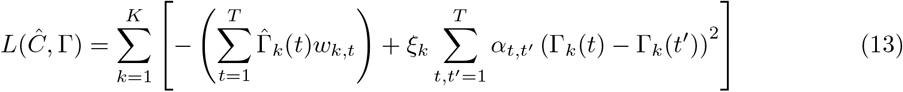

with constants

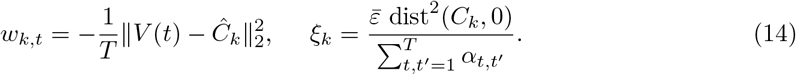 In the following, we simplify the objective function (13) into the standard QP form. Let us denote the diagonalization of matrix into vector

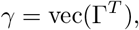

and introduce multi-index (*t, k*) of vector *γ* by

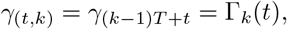

where *γ*_*j*_ is *j*-th component of *γ* ∈ ℝ^*KT*^. At first, notice that quadratic term

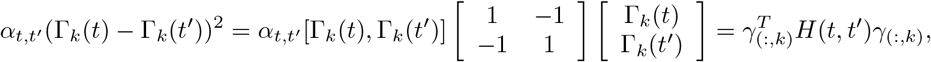

where *γ*_(:,*k*)_ = [*γ*_(1,*k*)_, … , *γ*_(*T,k*)_]^*T*^ ∈ ℝ^*T*^ and the components of matrix *H*(*t, t′*) ∈ ℝ^*T,T*^ are given by

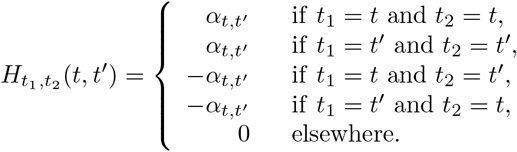 Using this notation, we are able to simplify the quadratic term of (13) into

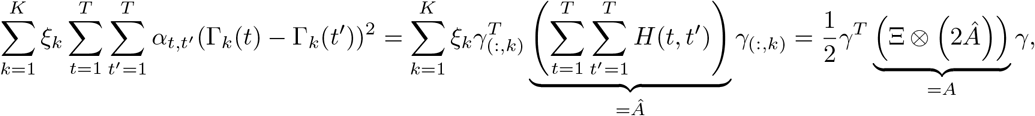

where Ξ = diag(*ξ*_1_, … , *ξ*_*K*_) ∈ ℝ^*K,K*^ is diagonal matrix and ⨂ denotes matrix Kronecker product. Matrix *A* ∈ ℝ^*KT,KT*^ is a block-diagonal matrix of *K* diagonal blocks 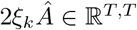. Let us remark that the matrix

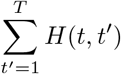

forms the Laplace matrix corresponding to graph of neighborhood of vortex *t* (stencil). Consequently, matrix 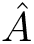 is composed from contributions from all stencils constructed in vortexes in the system. Such a matrix is symmetric positive semidefinite. The objective function can be written in the form of convex quadratic function

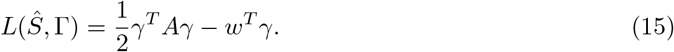 The feasible set (2) defines the lower bound constraints and equality constraints of the optimization problem. This feasible set is closed and convex, the objective function (15) is continuous, therefore using the Weierstrass Extreme value theorem [3], the optimization problem has always solution.
2. Before solving the QP problem (15), we assemble the Hessian matrix *A*, linear term *b*, and constraints Ω_Γ_ (2). The assembly of the linear term (14), where one has to sum *TK* values of distance functions between vectors of dimension *D*. If the complexity of chosen distance function evaluation is 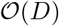, then the overall complexity is 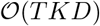.
3. Spectral Projected Gradient method for QP (SPG-QP, [7]) is an iterative algorithm for solving minimization problems of convex quadratic function 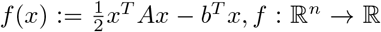 on closed convex feasible set Ω ⊂ ℝ^*n*^ defined by separable constraints with simple projections

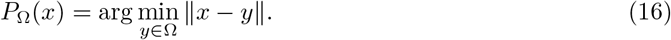 From the initial approximation *x*^0^ ∈ Ω, the process is generating the approximations *x*^it^ by

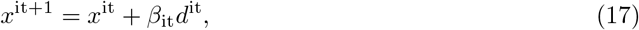

where *d*^it^ ∈ ℝ^*n*^ is *projected gradient* computed as

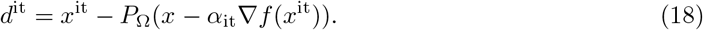 The step-size *α*_it_ is computed by Barzilai-Borwein rule [1]

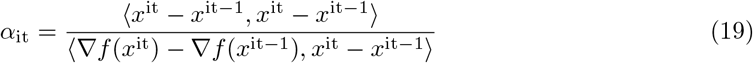

and the step-size *β*_it_ is a result of Grippo, Lampariello, and Lucidi line-search method [5] for satisfying so-called *generalized Armijo criteria*

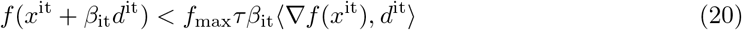

with safeguarding parameter *τ* ∈ (0, 1) and *f*_max_ is a maximum function value in previous *m* ≥ 1 iterations. The original SPG algorithm has been proposed by [2] for solving general optimization problems and the convergence is based on satisfaction of condition (20). Recently, [7] show that in the case of quadratic objective function, the line-search algorithm can be replaced by direct formula which satysfies (20)

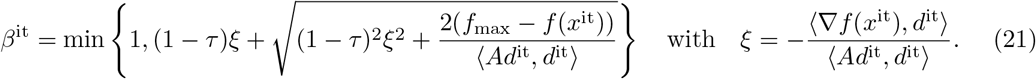 The algorithm non-monotonically decreases the norm of projected gradient and the function value until the stopping criteria is satisfied. In the general case, the most time-consuming operation is the multiplication by Hessian matrix *A*, all other computations includes the evaluation of scalar products. In our case, the matrix has a special pattern; it is a block-diagonal matrix of *K* diagonal blocks of band matrices. Computational complexity of multiplication with such a matrix is 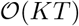. Please notice that the feasible set (2) is separable in *T* and the projection onto the set can be computed independently for each column of matrix Γ

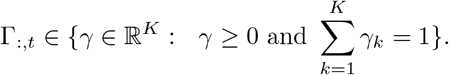 The projection onto each individual *simplex* is 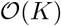, [4]. Summing up all the operations performed during one iteration of SPG-QP algorithm, the overall computation complexity is 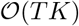.

#### 1. Theorem

*(The computational complexity of Algorithm 1)*

1. *Algorithm 1 generates the approximations with monotonically non-increasing objective function.*
2. *Let* dist *be Eulidean distance* (4). *One iteration of Algorithm 1 is* 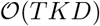.

*Proof.*

1. Since the iterations solves the optimization problems (6) and (7), we have

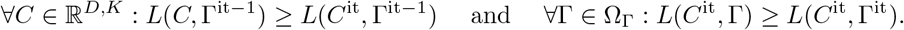 Choosing *C* = *C*^it−1^ and Γ = Γ^it−1^, we get

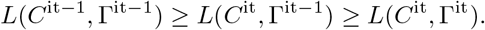
2. The statement is the consequence of Lemma 2 and Lemma 3.

### 2 Parallel Regularized Scalable Probabilistic Approximation Algorithm based on overlapping Domain Decomposition (DD-rSPA)

In the case of the computation in real-world applications, we are dealing with two main challenges: the computational demand (the number of operations that have to be performed to obtain the solution) and the memory limitation (the amount of information which can be processed by given machine). Both of these issues can be solved by High-Performance Computing (HPC). In this case, the algorithm runs on the machine which consists of several computational units (cores, processors, graphics processing unit) which are operating with distributed memory. The computational capacity of the largest supercomputers in the world can achieve more than 10^17^ FLOPS (floating-point operations per second) and can operate with several petabytes of memory. However, the massively parallel architectures cannot be efficiently utilized without appropriate massively parallel algorithms. For example in the case of discretized solution of partial differential equations with a huge number of variables, the original problem can be decomposed into smaller independent subproblems using so-called Domain Decomposition methods (DD). The idea is to solve subproblems in parallel using the individual computational units of the machine (i.e., nodes, cores, GPUs) and the only limitations arise in the case of global communication for the satisfaction of the continuity of global solution through domains. In practice, two different approaches are commonly used: overlapping DD, where the subdomains overlap by more than the interface (e.g., Schwarz alternating method or additive Schwarz method), and non-overlapping methods, where the subdomains intersect only on their interface (e.g., Balancing domain decomposition (BDDC), or Finite Element Tearing and Interconnecting (FETI)).

To analyze the problem of the global continuity and non-separability, suppose that we decompose the solution into two disjoint parts Γ_{1}_ and Γ_{2}_. Then the objective function of corresponding quadratic optimization Γ-problem (15) can be written as (after appropriate permutation of indexes)

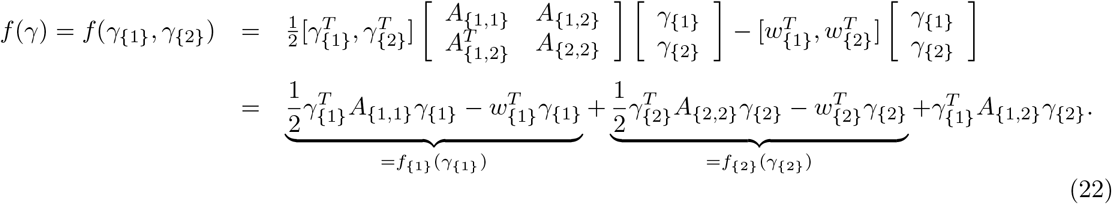

Using this equality, we can observe that the original minimization problem is separable into two disjoint minimization problems except the coupling term 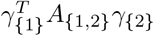. If we solve the problem separably to obtain *γ*_{1}_ and *γ*_{2}_ on separated computational units, we have to additionally handle with this term.

**Figure 1:**
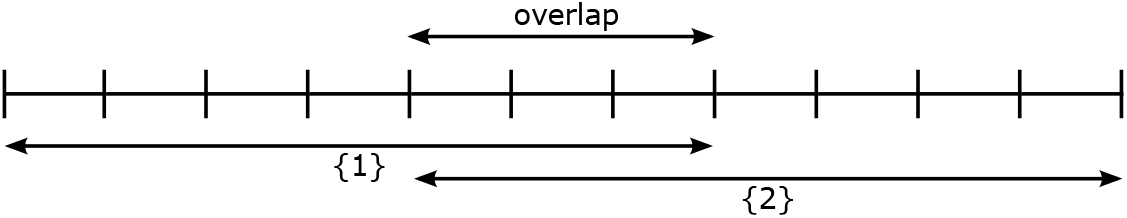
The overlapping Domain Decomposition: the domain is separated into continuous overlapping parts. Each computational unit computes the corresponding local solution, however, the information in overlap have to be communicated to satisfy the continuity of global solution through domains.

In our case, we implement the Schwartz Domain Decomposition method and separate the domain into overlapping domains, see Figure 1. For the demonstration, the figure presents the DD into two domains in 1D, but the approach is easily extendable to 3D and multiple domains in each direction, see Figure 2.

**Figure 2:**
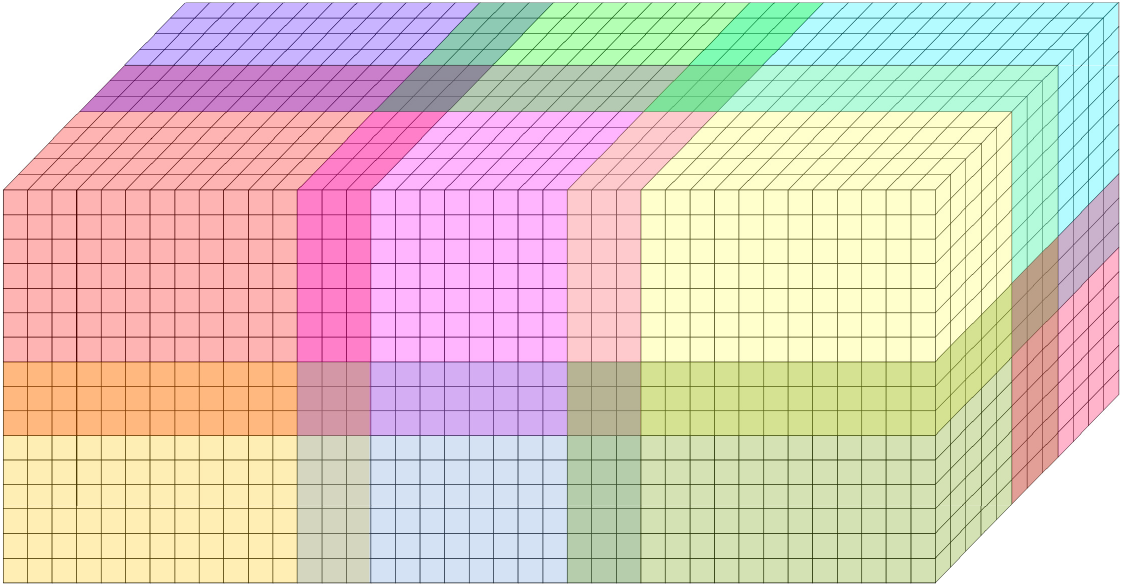
The overlapping Domain Decomposition in 3D: the simplest way how to decompose the 3D domain into domains is to introduce overlapping rectangular cuboids. Such a decomposition simlifies the implementation and follows the sparsity of Hessian matrix in Γ problem.

In the first step of algorithm, we solve the problem in each domain separately, i.e., we solve corresponding QP problem with appropriate block of the Hessian matrix and the block of linear term. Each domain sends the solution in overlap to the neighbouring domains and this vector is used for the computation of coupling term in local objective function. This operation can be written in terms of (22) - suppose that the local unknown part of the solution is *γ*_{*d*}_ and the rest of the solution is *γ*_{.}_. Then the objective function can be decomposed into (using the appropriate permutation of indexes)

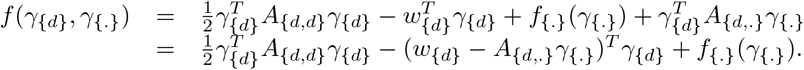

In the local domain, the unknown of the problem is the local *γ*_{*d*}_, therefore the term *f*_{.}_(*γ*_{.}_) is constant, does not have any impact on the optimizer, and can be ignored. Please notice that the coupling matrix *A*_{*d,.*}_ is a block of a sparse matrix and if the size of the overlap is sufficiently larger than the radius of the indicator function of the voxel neighborhood *α*_0_, then the overlap information from the neighboring domains is sufficient information for assembling the correct overall objective function. In the iterative process, we update the linear term using the information from neighbours, solve a new QP problem, and communicate the update of overlap to neighbours, see Algorithm 2. In each iteration, we compare the local solution in overlap with the solution obtained from neighbours and if the difference is sufficiently small, we stop the algorithm.

#### Algorithm 2: Schwartz Domain Decomposition method for solving Γ-problem in parallel.

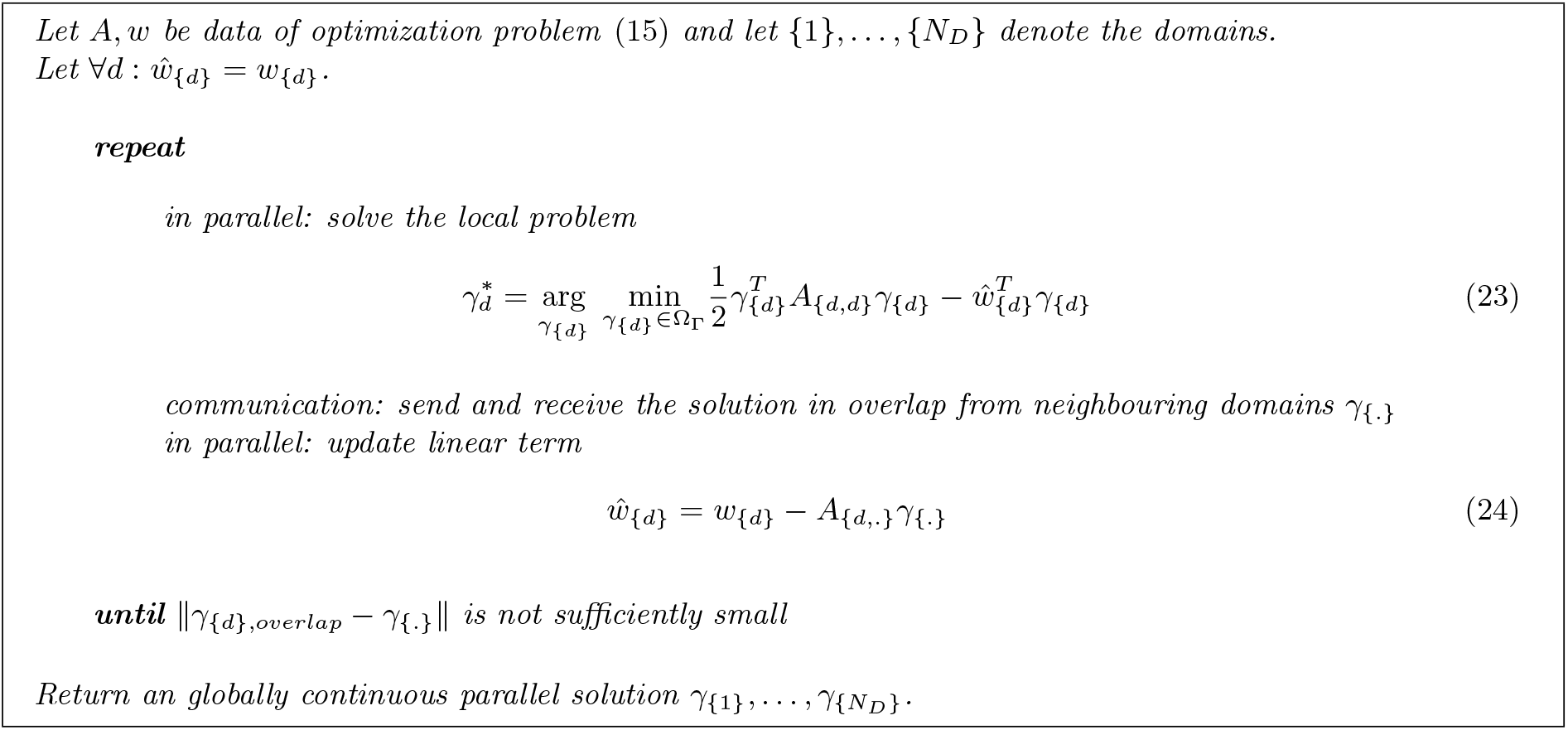

## Footnotes

1 Applying Ridge, Lasso and elastic net regularizations with respect to both variables *C* and Γ in problem (4) would result in regularization terms of the form +*ϵ*_*C*_∥*C*_*k*_∥ + *ϵ*_Γ_∥Γ_*k*_∥ and would require tuning at least the two regularization parameters *ϵ*_*C*_ and *ϵ*_Γ_.

